# Cortical representation of touch *in silico*

**DOI:** 10.1101/2020.11.06.371252

**Authors:** Chao Huang, Fleur Zeldenrust, Tansu Celikel

## Abstract

With its six layers and ~12000 neurons, a cortical column is a complex network whose function is plausibly greater than the sum of its constituents’. Functional characterization of its network components will require going beyond the brute-force modulation of the neural activity of a small group of neurons. Here we introduce an open-source, biologically inspired, computationally efficient network model of the somatosensory cortex’s granular and supragranular layers after reconstructing the barrel cortex in soma resolution. Comparisons of the network activity to empirical observations showed that the *in silico* network replicates the known properties of touch representations and whisker deprivation-induced changes in synaptic strength induced *in vivo*. Simulations show that the history of the membrane potential acts as a spatial filter that determines the presynaptic population of neurons contributing to a post-synaptic action potential; this spatial filtering might be critical for synaptic integration of top-down and bottom-up information.

## Introduction

One of the grand challenges in neuroscience is to mechanistically describe the cerebral cortical function. Numerous studies have identified the organizational principles of cortical circuits in various cortical areas across model systems by describing the principles of neuronal classification, cell-type specific projection patterns, input-output mapping across cortical layers, and by functional characterization of the anatomically identified neurons upon simple stimulation conditions, (see e.g. Douglas and Martin, 2004; Markram et al., 2015). Although a wiring-diagram approach is critical for a *structural* description of the network, relating the anatomical structure to network *function* will require a detailed study of the dynamical processes in single neurons as well as neural populations (Douglas and Martin, 2007; O’Connor et al., 2009). Or, in other words, one of the best ways to understand the functioning of the brain is trying to build one (Einevoll et al., 2019; Eliasmith and Trujillo, 2014). Accordingly, a large number of large-scale reconstructed computational models of cortical function (see Supplemental Table 1, the discussion section and this recent review (Fan and Markram, 2019)), including macaque (Chariker et al., 2016; Schmidt et al., 2018a, 2018b; Schuecker et al., 2017; Zhu et al., 2009), cat (Ananthanarayanan et al., 2009) and mouse/rat (Arkhipov et al., 2018; Billeh et al., 2019) visual cortex, rat auditory cortex (Traub et al., 2005), rat hindlimb sensory cortex (Markram et al., 2015), cerebellum (Sudhakar et al., 2017) and “stereotypical” mammalian neocortex (Izhikevich and Edelman, 2008; Markram, 2006; Potjans and Diesmann, 2014; Reimann et al., 2013; Tomsett et al., 2015), have been introduced, where neuronal dynamics are approximated using neuron models that range from integrate-and-fire point neurons (Ananthanarayanan et al. 2009, Sharp et al., 2014; Zhu et al., 2009, Potjans & Diesmann, 2014, Chariker et al. 2016, Bernardi et al. 2020, Schmidt et al., 2018a, Schmidt et a. 2018b, Schuecker et al. 2017) to morphologically reconstructed multi-compartment neurons (Traub et al. 2005, Markram et al. 2006, Izhikevich & Edelman 2008, Reimann et al. 2013, Markram et al. 2015, Tomsett et al., 2015, Sudhakar et al. 2017, Arkhipov et al. 2018, Billeh et al. 2019). These models have given insights in a range of topics including the nature of the local field potentials (Reimann et al., 2013; Tomsett et al., 2015), mechanisms of state transitions (Markram et al., 2015), frequency selectivity (Zhu et al., 2009), the influence of single-neuron properties on network activity (Arkhipov et al. 2018) and the relation between connectivity patterns and single-cell functional properties (i.e. receptive fields, Billeh et al. 2019).

With its topographical organization, well-characterized structural and functional organization, and its ever growing number of publicly available molecular, cellular and behavioural big datasets (Azarfar et al., 2018b; da Silva Lantyer et al., 2018; Kole et al., 2017, 2018a), the barrel column is ideally suited as a model system for computational reconstruction of circuit organization and function. Accordingly, large-scale computational models of the rodent barrel cortex ranging from detailed reconstructed models that need to be run on a supercomputer (Phoka et al., 2012; Sharp et al., 2014) to much less detailed and computationally expensive models (Bernardi et al., 2020) have been developed. However, currently, there are not any publicly available tools available for biologically realistic network modeling that can be performed using a standard computer of today. Therefore, we developed an open-source biologically constrained computational network model of the granular and supragranular layers of the barrel cortex along with the ventroposterior medial thalamus. It is a detailed model, with cortical cell densities based on the reconstructions in soma resolution presented herein and our previous work on a temporal variation in response dynamics (Huang et al., 2016). The code can be run on a desktop computer with or without a CUDA enabled GPU and is available for download on GitHub (https://github.com/DepartmentofNeurophysiology/Cortical-representation-of-touch-in-silico). Here we show that this barrel cortex *in silico* can predict (a) emergent whisker representations, (b) changes in the synaptic strength upon whisker deprivation, (c) network representation of touch from behavioral data, using only the information extracted from whisker tracking. The model will help novel principles of information processing (Huang et al., 2020).

## Results

### Anatomical organization of the barrel cortex

Just like most other neocortical areas, barrel columns consist of six layers with distinct molecular fingerprints and tens of different neural classes (Azarfar et al., 2018a; Fox, 2018; Kole et al., 2018b; Markram et al., 2004; Oberlaender et al., 2012; Thomson and Lamy, 2007). The reconstruction of the network in soma resolution (Figure 1, for detailed methods, see Materials and Methods) shows that the laminar distribution of cell-types varies significantly across layers. Similar to the laminar borders observed in the traditional Nissl staining, staining the column with neuronal nuclear antibody anti-NeuN, hereafter NeuN, results in a higher cellular density in Layer (L)4 and lower layers of L3 in comparison to L2 and L5-6. Inhibitory neurons stained with anti-GABA do not obey the laminar borders as outlined by the NeuN and display near equal densities in lower L4, L5b, and L1. Specific inhibitory neuron markers, however, have distinct expression patterns across the laminae: While Calretinin neurons are predominantly found in the L4/L3 border, Somatostatin neurons are preferentially located in the infragranular layers (Figure 1E). Parvalbumin-positive interneurons, on the other hand, are found at higher densities in L4 and L5. (Figure 1E). The cellular distributions in the canonical D-row column can be found in Supplemental Table 2.

**Figure 1.**
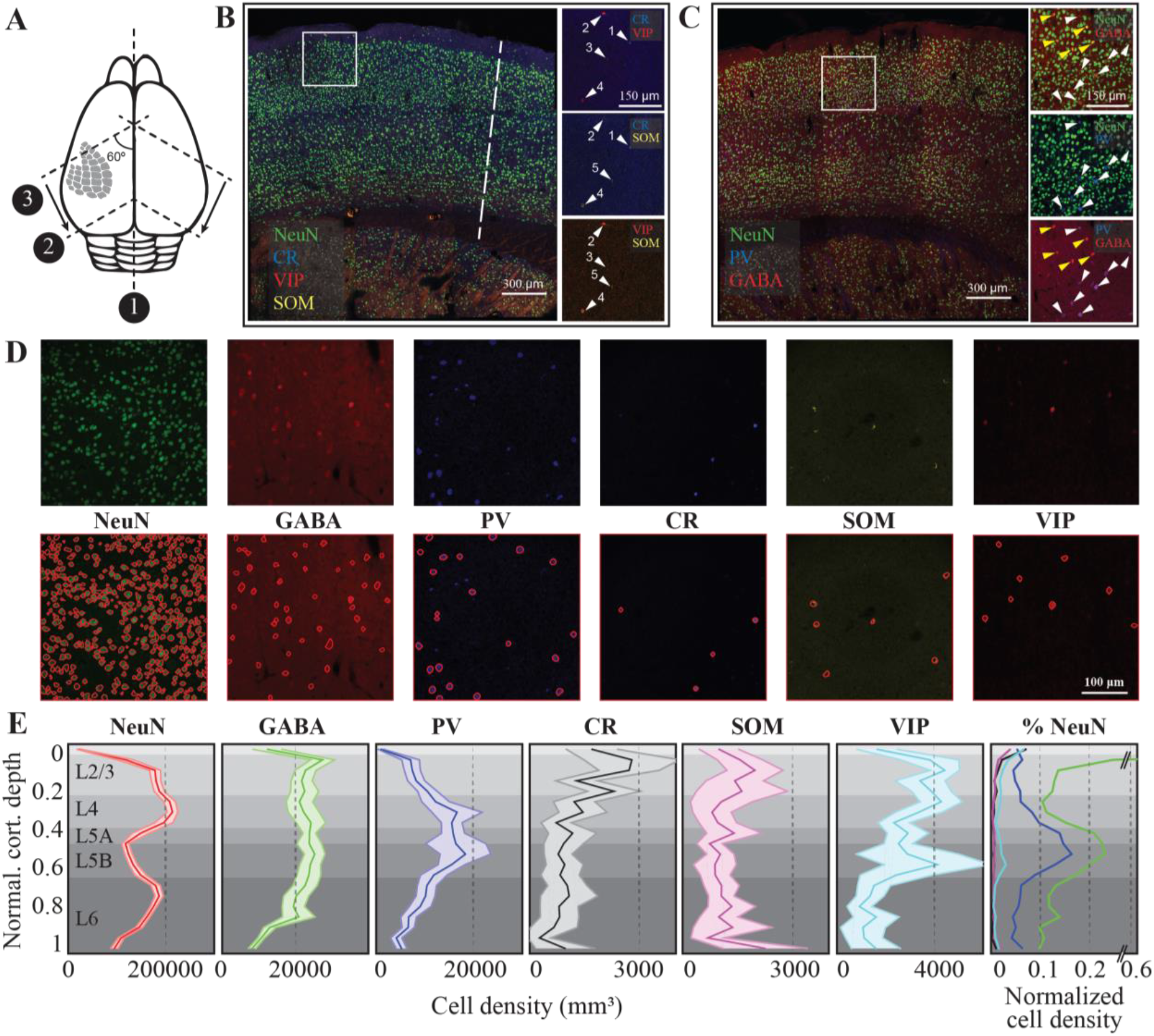
The anatomy of the canonical cortical column in the mouse barrel cortex. **(A)** Schematic representation of the slicing approach. Numbers refer to the order of incision (1,2) and sectioning (3) (see Materials and Methods for details). **(B-C)** Six monoclonal antibodies raised against select cellular markers were used for co-staining cellular classes. Insets show different staining patterns. Cell labeled with the same number is the same cell across different stainings. **(D)** Randomly selected raw images (top row) along with automatically detected cells in a 300×300×25 microm volume of fixed tissue (see Materials and Methods for details). **(E)** Density of identified cellular populations across the six cortical layers. The shaded regions represent 2 standard deviations from the mean (N=22 slices for NeuN, 12 for GABA, PV and 10 for CR, SOM, VIP; in average 3 columns in each slice from 3 animals, 5 hemispheres. Values are mean and std calculated from each slice). The last column represents the relative cellular density after normalizing the cell count to the number of NeuN positive neurons in a given layer.

### Stimulus representations in silico network

To create a network model, three components are necessary: 1) the distribution of the nodes, 2) edges and 3) a dynamic model of information transfer in single nodes. The first of these components, the distribution of nodes, was measured in the previous section (Figure 1). The second component, network connectivity, was determined using axonal and dendritic projection patterns (Egger et al., 2008; Feldmeyer et al., 2006, 2002; Helmstaedter et al., 2008; Lübke et al., 2003), which were approximated by 3-D Gaussian functions (see Materials and Methods and Supplemental Table 3), with the assumption that the probability that two neurons are connected is proportional to the degree of axonal-dendritic overlap between these two neurons (i.e Peter’s rule, (White, 1979)). For the third component, the dynamic model of single neurons, we modified the computationally efficient Izhikevich neuron model (Izhikevich, 2004, 2003) see Materials and Methods and Supplemental Table 4) to include the inverse relationship between the first derivative of the membrane potential, i.e the speed with which the synaptic depolarization rises, and the action potential threshold, so that the threshold is a function of the history of the membrane potential on (the membrane state (Huang et al., 2016; Zeldenrust et al., 2020)). This modification in the quadratic model did not affect the model’s ability to predict the timing of action potentials upon sustained current injection in soma (see Figure 2A; compare the middle column to (Izhikevich, 2004, 2003) and also correctly predicted the rate and timing changes associated with the membrane state at a single neuron resolution (Figure 2A).

**Figure 2.**
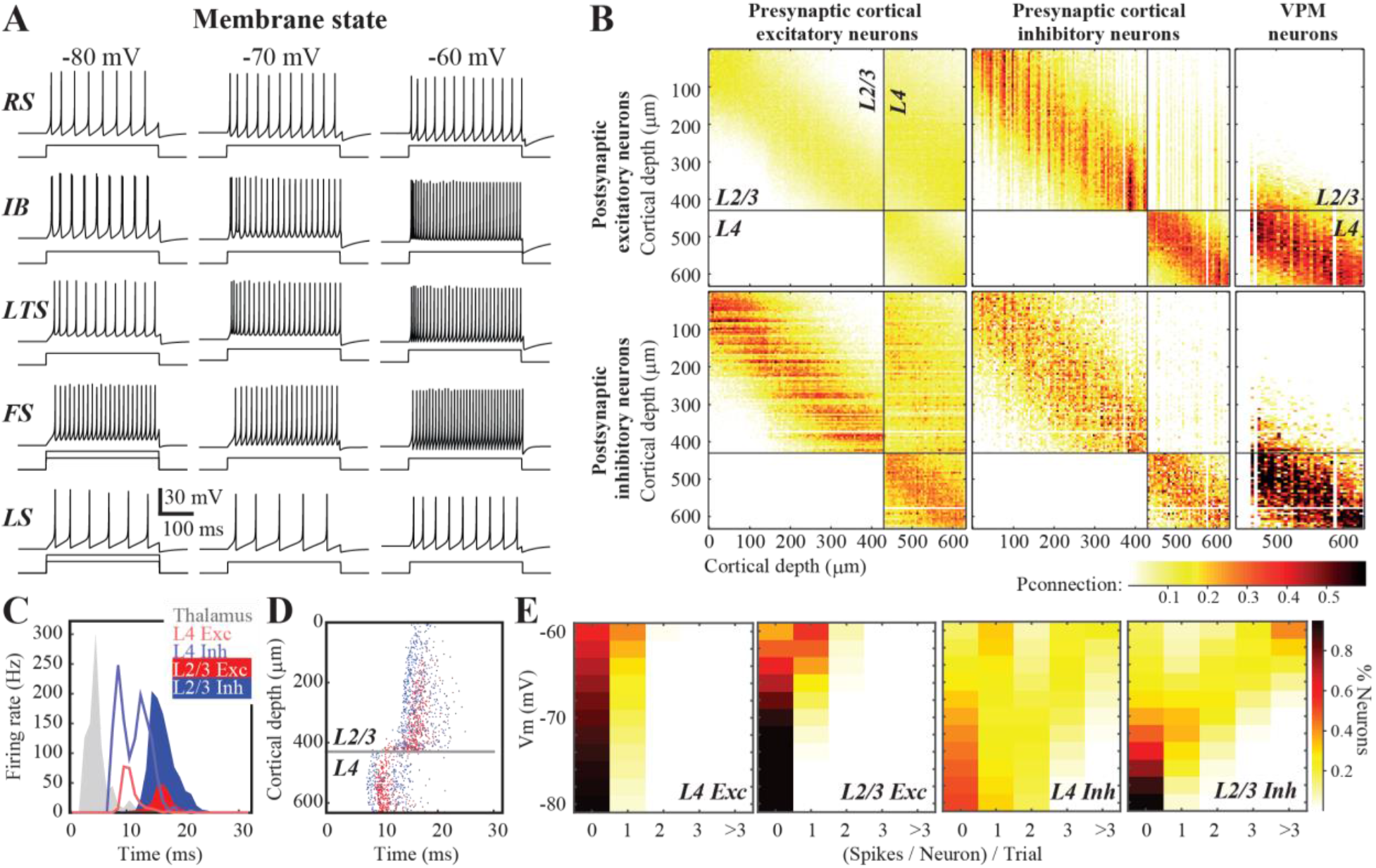
Neural activity and circuit connectivity *in silico*. **(A)** Spiking pattern of five electrically characterized cell classes upon somatic step-and-hold current injection across three membrane states. **(B)** The connectivity matrix across the network. **(C)** Emergent cortical activity upon thalamic stimulation, simulated as a response to a single whisker deflection (Petersen et al., 2008). Peristimulus time histograms (PSTHs) depict population responses across thalamus (modeled; see Materials and Methods for details), and cortical responses. **(D)** Same as in C but action potentials from neurons in the top 630 microm of the cortical column are shown. **(E)** Likelihood of spiking across identified neuron classes and membrane states.

With the completion of the three required components for functional network creation, we constructed a biologically constrained barrel cortical column *in silico*. Due to the general lack of experimental data on the pairwise connectivity between infragranular layer neurons and the rest of the network, in this version of the *in silico* column, we have constrained the network to the top 630 μm (Figure 2B), which is border between L4-L5 in the mouse. As the granular layer (L4) is the principal recipient of the thalamic inputs (Azarfar et al., 2018a) and strongly drives the supragranular (L1-3) layers, before the cross-columnar integration takes place across the upper L2/3, this model provides an *in silico* simulation environment for the first three stages of thalamocortical and intracortical information processing that involves supragranular and granular layers.

In the simulated network, stimulus-evoked activity spreads across the network from ventroposterior medial nucleus (VPM) to L2/3 with latencies comparable to those observed in biological networks under anesthesia (Figure 2C, (Allen et al., 2003; Armstrong-James et al., 1992; Celikel et al., 2004). Inhibitory neurons had an earlier onset of spiking with a peak latency of 8.2±0.6 ms (mean±std) in L4 (Figure 2C), which corresponds to <3 ms conduction delay, calculated from the population peristimulus time histograms (Figure 2C). These delays are similar to previous observations *in vivo* (Condylis et al., 2020; Dudai et al., 2020; Sermet et al., 2019; Swadlow, 2003, 1995). In terms of the latency to an action potential, neurons across the entire depth of L4 were homogenous with the exception that those closer to the L3 border showed a delayed spiking (Figure 2D). As the feed-forward projections originating from L4 are the main inputs to the L2/3 neurons, the activity *in silico* naturally follows the latency distribution observed *in vivo* across the cortical layers, with L2 neurons generating action potential up to 4 ms later than the lower L3 neurons (Celikel et al., 2004); Figure 2C). Independent from the actual location of the neuron within the *silico* network, however, inhibitory neurons have an earlier onset of spiking as compared to the neighboring excitatory neurons within the layer (Figure 2D).

The spiking probability varies significantly across layers and neuron types *in vivo* (Celikel et al., 2004; De Kock et al., 2007; Gentet et al., 2012, 2010; O’Connor et al., 2010) and *in silico* (Figure 2D). Excitatory neurons respond to the stimulus sparsely, as the probability of a given neuron to generate an action potential at a given trial is low. When the stimulus does yield a suprathreshold response, the neuron typically generates a single action potential (Figure 2E). The response probability and the number of action potentials/stimulus depend on the laminar location of the neuron, its cell type and its subthreshold membrane potential prior to the stimulus (Figure 2E; (Zeldenrust et al., 2020)) The laminar position of the neuron, be it excitatory or inhibitory, does not play a role in state-dependent changes in excitability at the single neuron level, although neurons in the supragranular layers respond on average more reliably to stimuli. The only exception to this rule is when the stimulus arrives in a hyperpolarized membrane state; if the resting membrane potential prior to the stimulus onset averaged <-75 mV, both excitatory and inhibitory neurons in L2/3 display failure rates higher than corresponding L4 neurons in the same membrane state (Figure 2E). This suggests that in hyperpolarized states, the activity of the supragranular layer is effectively uncoupled from the bottom-up sensory input.

### The source of response variability in silico

In a network where information propagates across synaptically coupled neurons via relatively weak, failure-prone and sparse connections, identical stimuli in the periphery will evoke distinct neural activation patterns, even if the measured spike rate and time are constant across presynaptic populations (given the stochasticity of the presynaptic population contributing to the postsynaptic spiking). Accordingly, neural representations in a biologically inspired *silico* network are expected to vary as a result of both the presynaptic spike timing variability and the changes in *effective connectivity* between layers and across trials discussed in the previous section.

To quantify the extent of the response variability *in silico*, we simulated the cortical responses to thalamic inputs in two conditions: (1) in every trial each thalamic spike train was generated as a result of an inhomogeneous Poisson process, constrained by the PSTH (see Figure 3A), or (2) a single realization of (1) was repeated over trials, so there was no trial-to-trial variability in the thalamic spike trains (see Figure 3B) and the thalamic spike trains were identical across trials. While the former condition creates variability in spike timing and the rate at the single thalamic neuron resolution, the latter condition preserves the rate and timing of the thalamic input onto the postsynaptic cortical neurons across trials. The results showed that the effective connectivity, i.e. which presynaptic neurons contribute to the firing of a postsynaptic neuron in a given trial, is a major contributor to the response variability (Figure 3). This contribution was independent of the membrane state of the postsynaptic neuron and the neuron class, although the variability increased with membrane depolarization (Figure 3, A2-A3).

**Figure 3.**
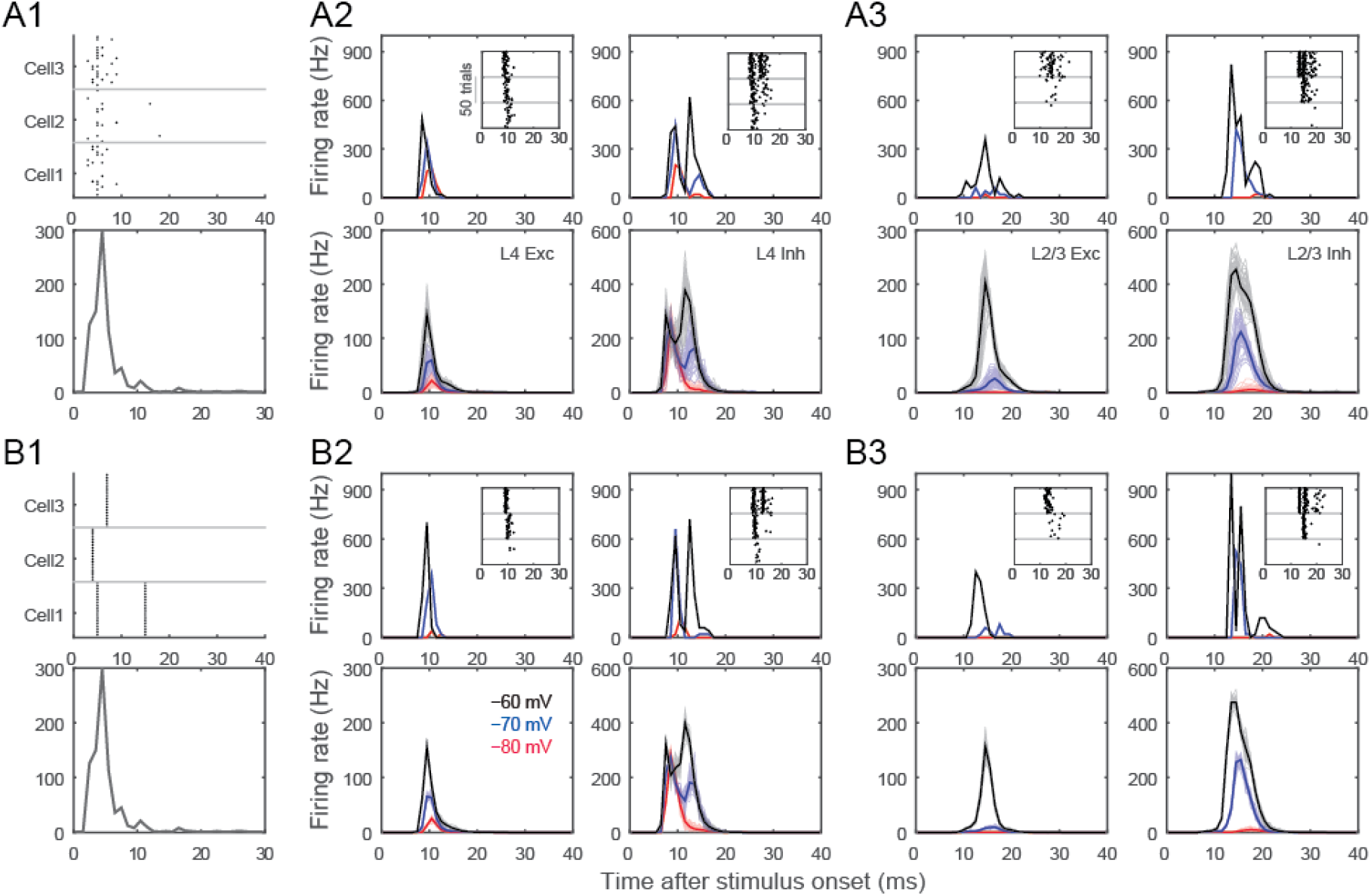
Variability of stimulus representations in silico. **(A1)** The population activity in thalamus is constrained by the PSTH as spike timing for individual cells are drawn according to Poisson-distributions. Raster plots exemplify the spiking responses of 3 representative thalamic neurons across 20 trials (upper panel) drawn from the population PSTH (lower panel). **(A2)** Representation of the thalamic input in single (upper panels) neurons and populations (lower panels). Left column: L4 excitatory, right column: L4 inhibitory neurons. PSTHs represent neural responses to 50 thalamic stimulation across three different membrane states (black: −60 mV, blue: −70 mV, red: −80 mV) **(A3)** Same as in A2, but for L2/3 excitatory and inhibitory neurons. **(B1)** The population PSTH in thalamus is the same as in A1 (lower panel), but spike timing and rate of individual thalamic neurons’ spiking is constant across trials (see raster plots in upper panel). **(B2, B3)** same as in A2, A3, but show cortical response to stereotypic thalamic inputs. Note that even when the same thalamic input pattern was used to stimulate the network, neurons still showed spike timing variability due to synaptic failures and synaptic strength variations although the variance was greatly reduced as the effective connectivity in the network was kept constant.

### Stimulus representations in L4 in silico

Thalamic neurons project extensively to cortical L4, and diffusely to the L3/L4 and L5b/6 borders (Arnold et al., 2001; Oberlaender et al., 2012; Sermet et al., 2019). This thalamocortical input is the principal pathway that carries the feedforward excitatory drive, carrying the bottom-up sensory information (Azarfar et al., 2018a) L4 representations of the sensory input are characterized by sparse neural representations *in vivo* (Aguilar, 2005; Celikel et al., 2004; De Kock et al., 2007) and *in silico* (Figure 4). Thalamic input modeling the principal whisker’s stimulation *in vivo* results in a significant firing rate modulation (two orders of magnitude, between 0.02-2.2 spikes/stimulus/cell) in the network, depending on the membrane states of the L4 neurons prior to the stimulus arrival as well as the neuronal class studied (at *vr=-80* mV, excitatory neurons fire at 0.06±0.11 spikes/stimulus, range 0-0.82; inhibitory neurons, 0.68±0.71 spikes/stimulus, range 0-2.22; at *vr*=-60mV, excitatory neurons, 0.44±0.30 spikes/stimulus, range 0-1.96; inhibitory neurons, 2.13±1.48 spikes/stimulus, range 0.02-6.54; values show mean±std). While excitatory neurons fire sparsely, inhibitory neurons spike with higher reliability (Figure 4C). The resting membrane potential changes the properties of excitatory neurons firing, as L4 excitatory neurons switch from a sparse representation (i.e. the probability of spiking for each neuron per stimulus is low, and when neurons spike they typically fire single action potentials) to less sparse spiking as membrane potential depolarizes (Figure 4E). The inhibitory neural population, on the other hand, undergoes rate scaling as the resting membrane potential is depolarized (Figure 4E). Hence for the neural coding of stimuli in L4, the membrane state acts as a state-switch for excitatory neurons and a gain-modulator for the inhibitory neurons in the principal whisker’s cortical column.

**Figure 4.**
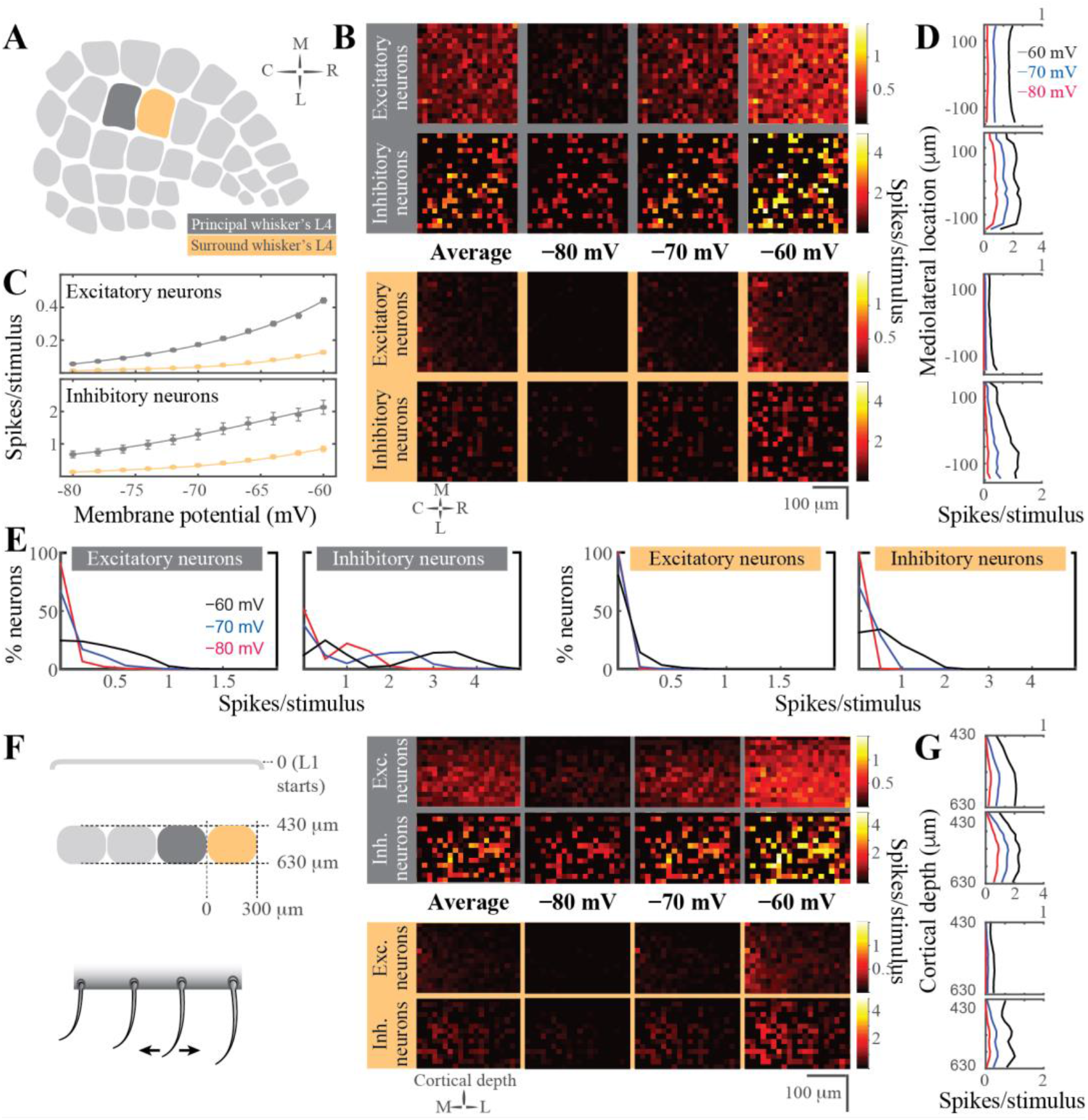
Stimulus evoked representations in cortical layer 4 *in silico*. **(A)** Schematic representation of the spatial orientation of the simulated network. The visualizations are in the tangential plane. The principal cortical column is the D2 whisker’s column. **(B)** Average neuronal response in rostro-caudal (RC) and medio-lateral (ML) planes, across different resting membrane states (pixel size 15×15 μm in cortical tissue). The figurines on the grey shaded background display the response in the principal whisker’s cortical column; the yellow background shows the activity in the first order surrounding L4. **(C)** Average firing rate of excitatory (top) and inhibitory neurons (bottom) in the network as a function of the resting membrane potential before stimulus onset in the principal (top) and surround (bottom) whisker’s L4. **(D)** Average firing rate in the ML axis across the membrane states. **(E)** Distribution of the spiking response per stimulus across neuron classes and membrane states. **(F)** Left: Schematic representation of the coronal orientation of the visualized network. Right: Average neuronal response across the dorsoventral plane in L4 (pixel size 15×15 μm in cortical tissue). **(G)** Average firing rate across cortical depth.

The spatial distribution of synaptic inputs in a network is primarily constrained by the axo-dendritic overlap across the synaptically connected neurons. Accordingly, with diffuse axonal projections of thalamic neurons, and spatially constrained dendritic branching to the barrel borders, excitatory and inhibitory L4 neurons along the rostro-caudal (RC) and medio-lateral (MC) planes do not display a spatial bias in the tangential plane (Figure 4B). Unlike this spatial homogeneity of L4 responses to the stimulus, preferential laminar targeting of the thalamic input results in a higher likelihood of spiking in the bottom portion of the barrel, especially for postsynaptic excitatory neurons (Figure 4F).

The topographical nature of the representation of whisker touch dictates that each neuron has a preferred whisker, called the principal whisker, which evokes the largest number of action potentials upon deflection (Brecht and Sakmann, 2002; Foeller et al., 2005). However the receptive fields of cortical neurons are rarely (if ever) constrained to a single whisker, as multi-whisker receptive fields in the thalamus (Aguilar, 2005; Armstrong-James and Callahan, 1991; Diamond et al., 1992; Kwegyir-Afful et al., 2005; Simons and Carvell, 1989) and cross-columnar projections in the cortex (Egger et al., 2008) ensure that each neuron receives information from multiple whiskers. Responses to the surround whiskers are always weaker, in number of spikes per stimulus, and arrive with a delay compared to the principal whisker deflection (Brecht and Sakmann, 2002). This relationship is preserved *in silico* representations of touch presented here (Figure 4B, C, F). Principal *vs* surround whiskers activate excitatory and inhibitory neurons similarly, although evoked representations of surround whiskers are invariably weaker (Figure 4B). Similar to the principal whisker deflection, surround whisker stimulation results in largely homogenous representations across the RC-ML axis (Figure 4B) even if the postsynaptic spiking is constrained to depolarized membrane states. The sublaminar activation pattern in L4 results in a higher likelihood of spiking in the bottom half of L4, even after surround whisker stimulation (Figure 4F).

One main difference between the principal vs surround representations is the role of the membrane state in the modulation of network activity. Unlike the differential role of the resting membrane potential in encoding principal whisker touch across the excitatory and inhibitory networks, the contribution of the different membrane states to surround whisker representation slowly (but predictably) varies across different membrane states (Figure 4C). Most excitatory and inhibitory neurons in the surround L4 do not represent the stimulus information during the quiescent hyperpolarized membrane state, resulting in principal whisker specific cortical representations. In the depolarized membrane states, the probability of spiking disproportionately increases for the inhibitory neurons.

### Stimulus representations in the supragranular layers in silico

Feedforward L4 projections are powerful modulators of supragranular layers and bring the bottom-up information from the sensory periphery for eventual cross-columnar integration primarily via L2, and less so via upper L3 neurons (Kerr et al., 2007; Petersen, 2007; Petersen and Sakmann, 2001). Principles of sensory representations by L2/3 *in silico* (Figure 5) are generally similar to the L4 neurons, with the exceptions that (1) supragranular excitatory neurons have an increased probability of firing during surround whisker stimulation, and (2) the spatial localization of a neuron has predictive power for its response properties.

**Figure 5.**
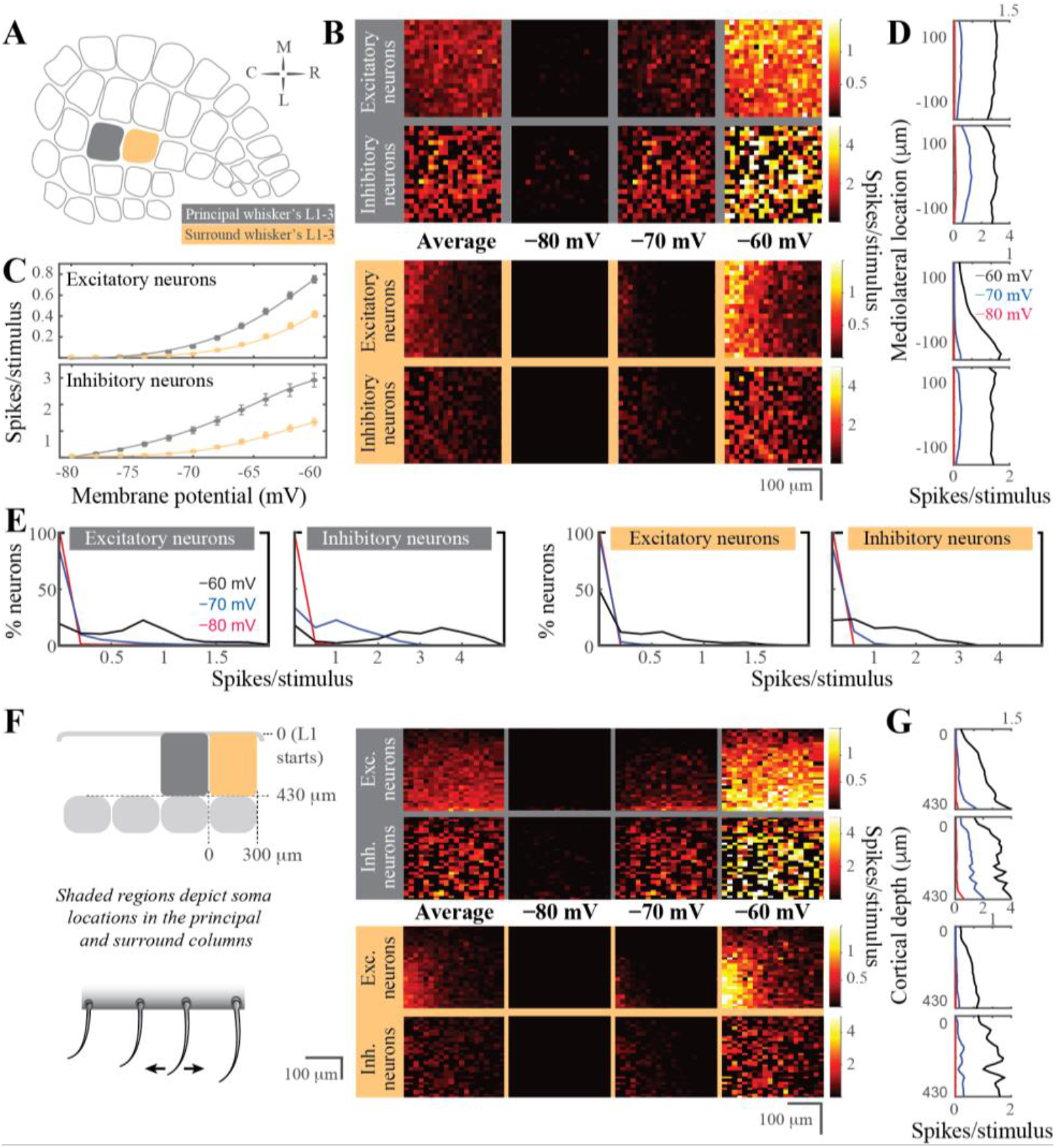
Stimulus evoked representations in the supragranular layers of the barrel cortical network *in silico*. **(A)** Schematic representation of the spatial orientation of the simulated network in the tangential plane. The principal cortical column is the D2 whisker’s column. **(B)** Average neuronal response mapped onto rostro-caudal (RC) and medio-lateral (ML) planes, across resting membrane states (pixel size 15×15 μm in cortical tissue). The figurines on the grey shaded background display the response in the principal whisker’s cortical column; yellow background shows the activity in the first order surrounding supragranular layers. **(C)** Average firing rate of excitatory (top) and inhibitory neurons (bottom) in the network as a function of the resting membrane potential before stimulus onset in the principal (top) and surround (bottom) whisker’s cortical network. **(D)** Average firing rate in the ML axis across the membrane states. **(E)** Distribution of the spiking response per stimulus across neuron classes and membrane states. **(F)** Left: Schematic representation of the coronal orientation of the visualized network. Right: Average neuronal response across the dorsoventral plane in L4 (pixel size 15×15 μm in cortical tissue). **(G)** Average firing rate across cortical depth.

Unlike the granular layer representations of the stimulus in the quiescent membrane states, L2/3 excitatory neurons are completely silent at hyperpolarized membrane potentials, suggesting that the bottom-up thalamocortical information is decoupled from the rest of the cortical circuits that originate from the supragranular layers. The lack of spiking is not specific to the excitatory neurons, inhibitory neurons are similarly unresponsive to the L4 input if the resting membrane potential was hyperpolarized (Figure 5C). Although inhibitory neurons fire stimulus-evoked action potentials at hyperpolarized membrane potentials (< −70 mV), the net effect of the membrane potential on suppressing cortical propagation of information via L2 is maintained across both classes of neurons (Figure 5). The lack of stimulus-evoked spiking in the surround column Figure 5 in resting membrane potentials < −70 mV and the changes in the spike probability described before suggest that sensory representations are weak but specific to the principal whisker column during the quiescent states *in vivo*.

Given that the neuronal excitability changes with the membrane state, that the neural thresholds depend on the stimulus and membrane potential history and that each neuron will (not necessarily linearly) sum its inputs until this variable threshold, the effective connectivity within the network should change with the membrane state of the postsynaptic neuron. To visualize the effective connectivity we spatially mapped the presynaptic neurons that fired action potential(s) prior to the spiking of a postsynaptic neuron (Figure 6). As expected, the effective connectivity varied with the membrane state. With an increasing probability of L2/3 spiking in the depolarized membrane states, the contribution of the intralaminar input to the spiking increased, suggesting that in the depolarized membrane states, sensory representations are a function of feed-forward drive originating from L4 and local changes in excitability in L2/3. The latter component is likely to be modulated by top-down modulations as the state of the animal changes during, for example, active sensing, providing a mechanistic model how the bottom-up sensory information can be integrated with the top-down neuromodulatory influences.

**Figure 6.**
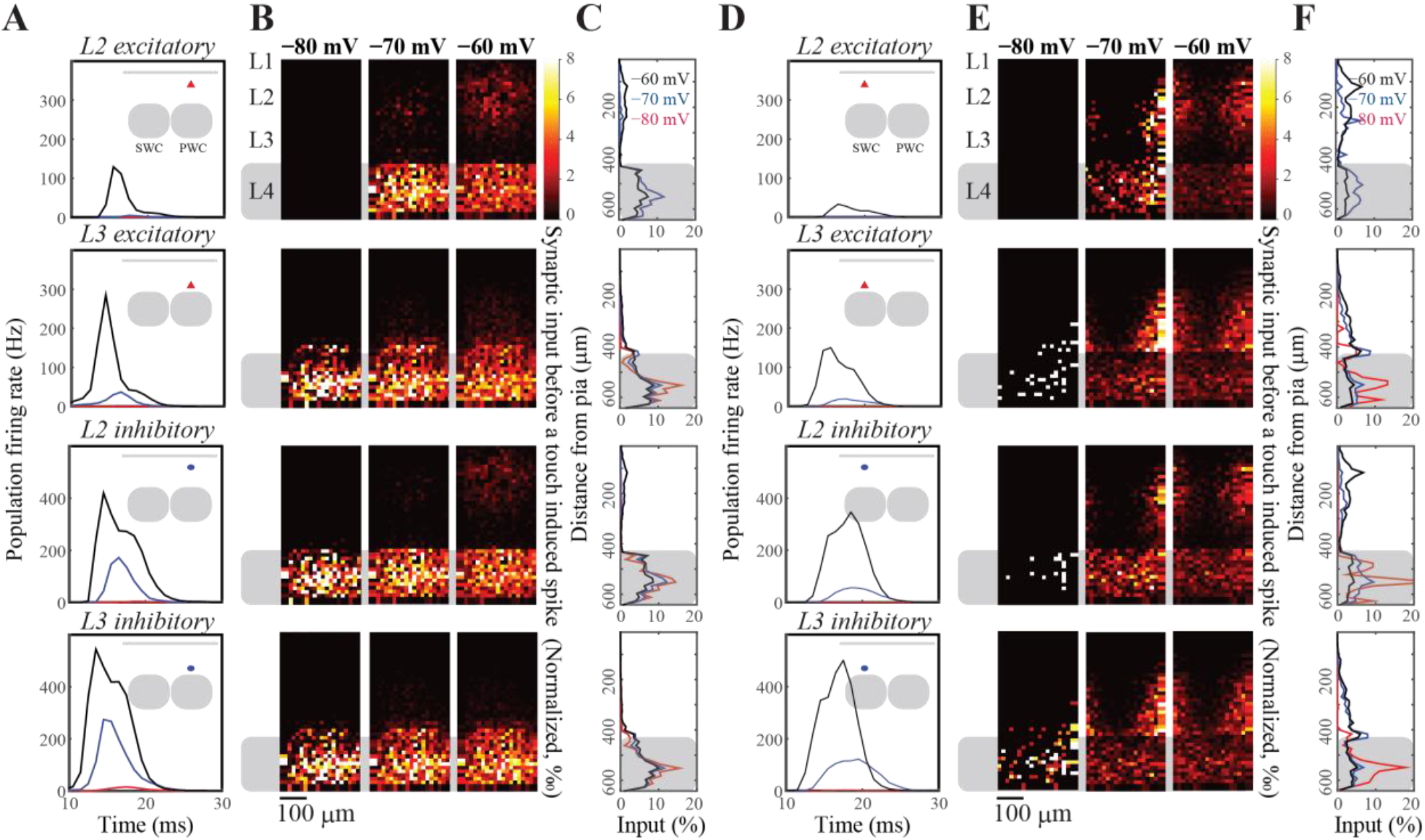
Visualization of the presynaptic population contributing to a postsynaptic action potential. We spatially mapped the neural activity across the granular and supragranular layers prior to an action potential in a given layer. The maps were averaged across all postsynaptic neurons that fire evoked action potentials during the simulations. (A) Population activity that drives L2 excitatory (first row), inhibitory (second row), L3 excitatory (third row) and L3 inhibitory (last row) neurons to spike in response to thalamic input. The first spike fired by aforementioned L2 or L3 neurons was used as the trigger to calculate the spike-triggered input map. Insert: schematic representation of the location of different cell populations in the barrel column. (B) Spike triggered spatial averaging (rows as in A); columns denote network activity observed across different resting membrane potentials. (C) Average depth distribution of excitatory inputs to drive a spike (rows as in A). (D-F) Same as A-C, but using surround whisker stimulation (SWC) instead of principal whisker stimulation (PWC).

### Experience-dependent plasticity of synaptic strength in silico

Neurons in the barrel cortex adapt to changes in sensory organ output as cortical circuits undergo plastic changes upon altered sensory input statistics (Allen et al., 2003; Clem et al., 2008; Feldman and Brecht, 2005; Kole et al., 2018b). These adaptive changes have long-lasting consequences in neural representations of touch. We have, therefore, integrated a spike-timing-dependent plasticity learning rule (Celikel et al., 2004) to enable plastic changes in neural representations of touch in silico. Figure 7 shows the implementation of the model on a 3-column model of the barrel cortex, layers 2-4 (Figure 7A). Each column receives its major synaptic input from its own respective whisker in the form of thalamic representations of whisker touch (see above), with the exception that the center column lacks a principal whisker, mimicking the whisker deprivation condition (Figure 7B).

**Figure 7.**
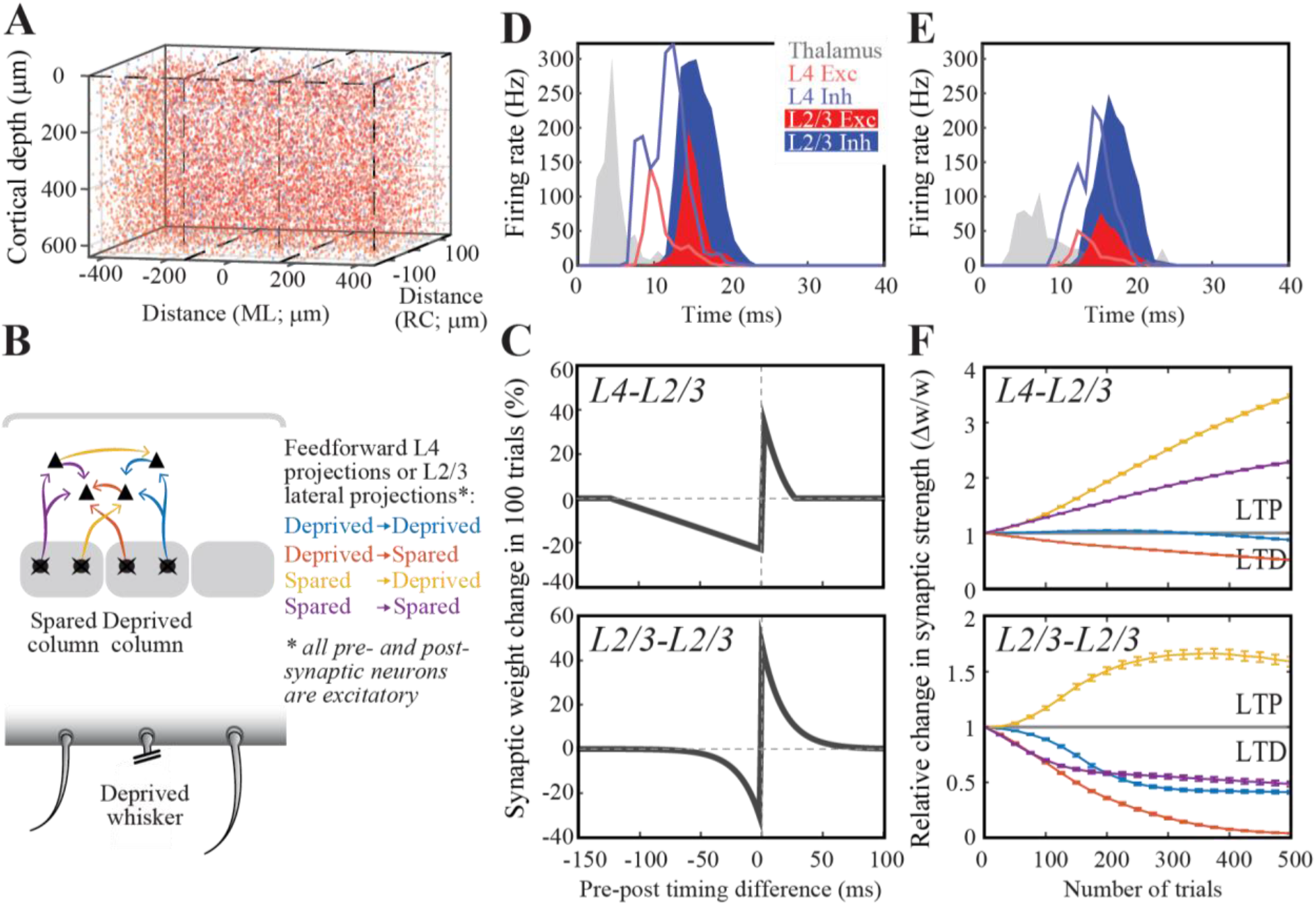
Spike-timing dependent map plasticity *in silico*. **(A)** A network model with 3 barrels. Cells in each column are randomly generated using distributions quantified in Figure 1. **(B)** Schematic representation of the feed-forward and intracolumnar networks in the upper layers of the somatosensory cortex. **(C)** Experimentally observed STDP learning rule in L4-L2/3 projections (top; Celikel et al 2004; see Materials and Methods) and for L2/3-L2/3 connections (bottom; Banerjee et al 2014). **(D)** Population PSTH for the spared columns, i.e most medial and most lateral columns in A1. **(E)** Population PSTH for the deprived, i.e. center, column. **(F)** Change in synaptic efficacy as a function of whisker deprivation in the simulated network. Color codes denote the whisker deprivation status of pre- and postsynaptic neurons’ location. Note that presynaptic neurons are always located in L4.

Employing empirically observed STDP rules in synapses at the feed-forward projections originating from L4 (Figure 7C; bottom) and the intracolumnar projections of L2/3 (Figure 7C; top) resulted in a reorganization of touch representation already within 100 trials, in agreement with the experimental observations in barrel cortical slices (Allen et al., 2003; Celikel et al., 2004). The model correctly predicted all the known pathways that are modified upon whisker deprivation including the potentiation in the spared whiskers’ L4-L2/3 projections (Clem et al., 2008), slow depression in the deprived cortical column’s L4-L2/3 projections (Bender et al., 2006) and plasticity of the oblique projections from L4 onto the neighboring L2/3 (Hardingham et al., 2011). The model further predicted a number of circuit changes, including the bidirectional changes across the cross-columnar projections between the spared and deprived columns, which could potentially explain the topographic map reorganization by receptive field plasticity

### Network representation of touch in vivo

As a final test of our in silico cortical column, we let it respond to an *in vivo*-like stimulation (Figure 8): as input to the network, we used recorded whisker angle (black) and curvature (red) from a freely moving rat in a pole localization task (data from (Peron et al., 2015)) made available as ‘ssc-2’ on CRCNS.org). We modeled thalamus as a network of 3 barreloids, each containing 200 ‘filter-and-fire’ neurons that respond to whisker angle, curvature, or a combination of both. The center barreloid was considered to be the principal barreloid for the spared whisker, whereas the other two were considered surround barreloids, with reduced probability (30% of original amplitude) and delayed (2.5 ms) response latency (Brecht et al., 2003; Brecht and Sakmann, 2002). The response of the network is tightly localized, both in time and place (Figure 8C,D). The network response is also quite sparse (Figure 8B,E), with each neuron firing at most a few spikes per trial. This response is a bit more sparse than typically observed (Peron et al., 2015), probably due to the lack of motor and top-down input in this model.

**Figure 8.**
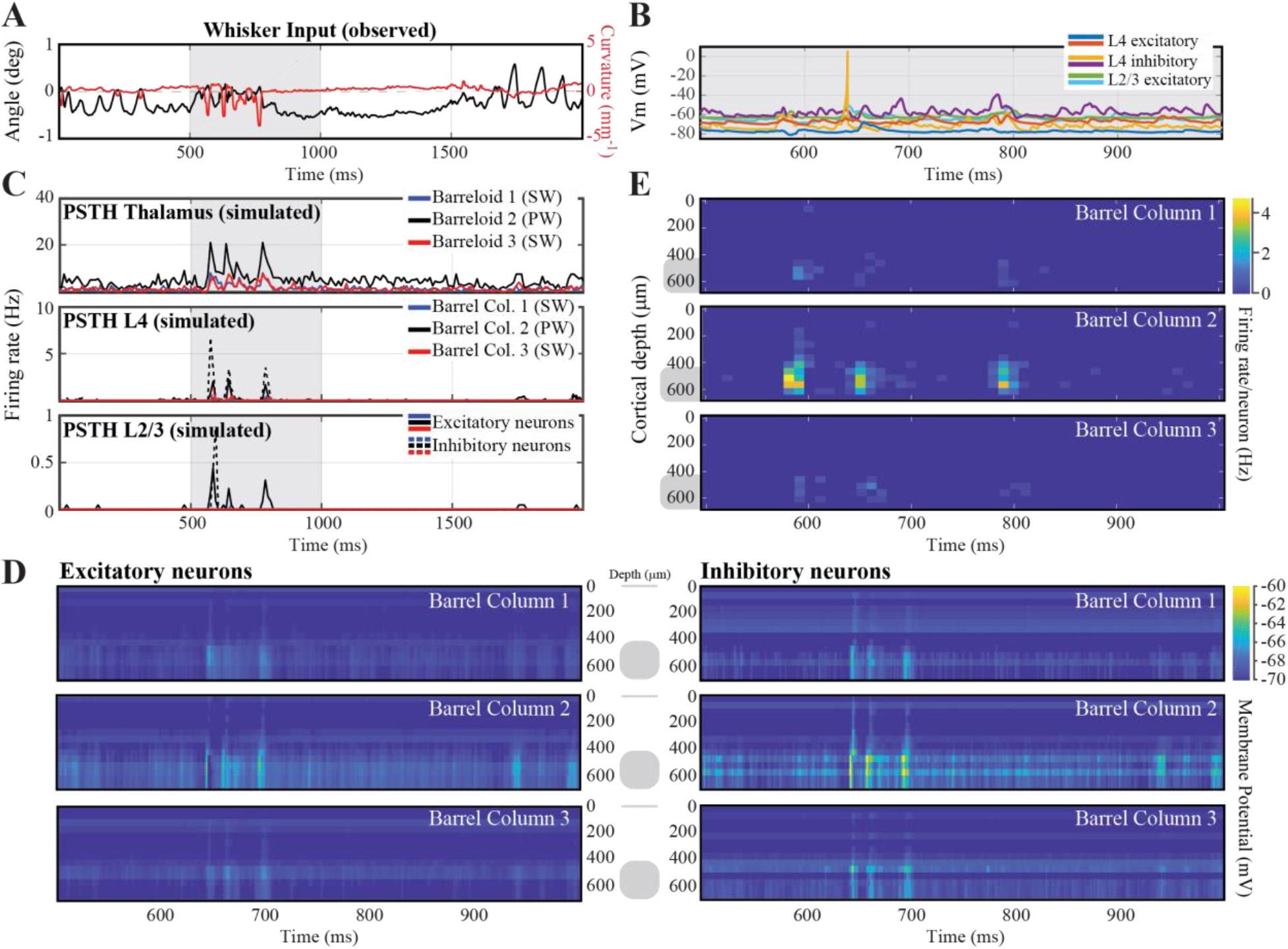
Network response to *in vivo*-like stimulation. **(A)** Input to the network: whisker angle (black) and curvature (red) from a freely moving rat in a pole localization task (data from (Peron et al., 2015), made available as ‘ssc-2’ on CRCNS.org). **(B)** Example voltage trace responses of 6 randomly chosen model neurons. **(C)** Peri-Stimulus Time Histograms (PSTHs) of the model-thalamus (top), L4 (middle) and L2/3 (bottom). The thalamus consists of 3 barreloids, each containing 200 ‘filter-and-fire’ neurons that respond to whisker angle, curvature or a combination of both. The central barreloid (black, 2) receives a stronger input, as this is the ‘stimulated’ barrel for the only spared whisker. Spike trains of the thalamus are sent to the cortical network model of L4 (middle), which sends its spike trains to L2/3 (bottom). These similarly consist of 3 barrels, of which the central (black, 2) barrel belongs to the spared whisker. **(D)** Average membrane potential of the excitatory (left) and inhibitory (right) model neurons as a function of cortical depth. L4 (barrel cortex) is denoted with a grey shaded shape. **(E)** Average firing rates of the model neurons as a function of cortical depth.

We compare the activity of a single barrel with evoked responses visualized using 2-photon imaging of calcium dynamics (Vogelstein et al., 2009). Although making a neuron-by-neuron comparison between networks is impossible, we can compare the overall activity of the networks. In both the recorded and the simulated networks, the activity is extremely sparse. The simulated network appears to have a few more neurons with a high firing frequency (Figure 9G), however, these do not adapt their firing frequency upon touch (Figure 9H), so they probably do not represent touch information (Peron et al., 2020). Otherwise, both networks show a comparable overall activity pattern.

**Figure 9.**
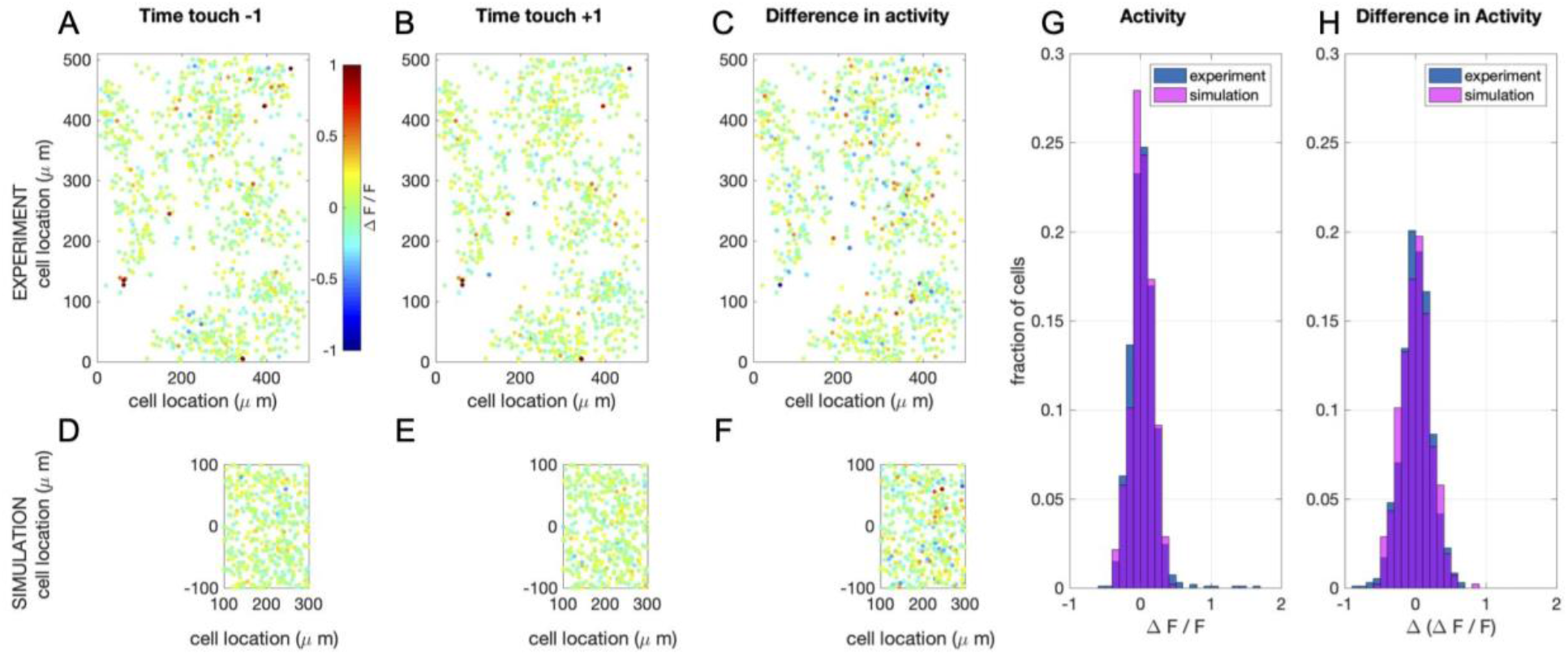
Simulation of calcium imaging experiment in L2/3. **(A)** Recorded (Peron et al., 2015) network response one (time) frame before touch (sampling frequency: 7 Hz; recorded volume: 6). **(B)** Recorded network response one frame after touch **(C)** Difference in network response between before and after touch. **(D-F)** Same as in A-C, but now for simulations (full simulation: single barrel including L23 (shown here) and L4 (see supplemental Figure 4). The fluorescence signal was calculated from network response following the method in (Vogelstein et al., 2009). Note that a recorded volume is larger than a single barrel. The frames are scaled accordingly. **(G)** Comparison of the distribution of activity of one frame after touch between the recorded and the simulated network. **(H)** Comparison of the distribution of the difference in activity between one frame before and after touch between the recorded and the simulated network.

## Discussion

Understanding the circuit mechanisms of touch will require studying the somatosensory cortex as a dynamical complex system. Given that the majority of research in the barrel system has thus far focused on the identification of circuit components the development of a computational model of the barrel cortex is not only necessary but also feasible. Accordingly, we here employed a three-tiered approach to (1) reconstruct the barrel cortex in soma resolution, (2) implement a model neuron whose spiking is a function of the network activity impinging onto postsynaptic neurons, and (3) axo-dendritically connect neurons in the column based on Peter’s rule and experimentally observed pairwise network connectivity (see Materials and Methods). We finally performed simulations in this network to compare neural representations of touch *in silico* to experimental observations from biological networks *in vivo*. As extensively discussed in the Results section, the simulations faithfully replicate experimental observations *in vivo* with high accuracy including, but not limited to, emergence of whisker representations, experience-dependent changes in synaptic strength and circuit representation of touch from behavioral data, using information from whisker displacement during tactile exploration. Thus, here we will focus on the methodological limitations and technical constraints of the network modeling as performed herein.

### Technical considerations for anatomical reconstruction of a stereotypical barrel column

One of the essential steps towards building a biologically plausible *silico* model of the mouse barrel cortex is to obtain the distribution patterns of different neuron types throughout the barrel cortex. In the current study, we directly visualized these distributions by labeling different types of neurons using cell-type specific markers and digitized the data using confocal scanning microscopy to ultimately reconstruct the cortex in soma resolution upon automated counting of all neurons, independent from whether the markers are nuclear or cytoplasmic. The identities of individual barrels in L4 can be reliably recognized based on GAD67 immunostaining (Supplemental Figure 3). However, due to difficulties in aligning images across consecutive sections, we could not consistently follow every barrel column across the entire cortical depth. Thus, in the current study, we only report average cell densities across a canonical barrel cortex rather than reconstructing the barrel cortex while preserving the columnar identity. Similarly, the *in silico* model places neurons and synapses stochastically every time a network is reconstructed, reflecting this inherent uncertainty. The advantage of this is, that simulations can be repeated over different realizations of networks with a similar structure, and this way it can be tested whether results are a general property of such networks or just a coincidental result of a particular realization of the network. It should be noted that, in the rat barrel cortex, the cell density across different barrel columns has been shown to be relatively constant (Meyer et al., 2013), making our density estimation likely to be accurate, as we employed a normalized volume for the entire column. Obviously, however, the absolute cell number in one barrel column could vary depending on the exact location of the barrel within the barrel cortex (Meyer et al., 2013).

Our automatic cell counting algorithm for nuclear cell counts is functionally similar to that employed in (Oberlaender et al., 2009). Compared to their method, we used lower threshold values to separate foreground objects from their background in order to capture weakly stained cells. This comes at the expense of an increased number of connected clusters. We thus employed more sophisticated methods to separate clusters of connected cells, based on both intensity and shape information, rather than simply assuming that there exists a single dominant cell population based on volume, which could lead to bias when the assumption is not met (Oberlaender et al., 2009). Our method does not require manual correction, and the counting results are comparable with manual counts (Supplemental Table 5). Furthermore, we also developed algorithms to enable source localization for the cytoplasmic signals, which allowed us to quantify cellular classes, like somatostatin neurons, that are characterized by non-nuclear markers. Together these approaches have resulted in the most detailed quantification of the network, going beyond the two-neuron group (i.e. excitatory vs inhibitory) clustering available in the literature.

Tissue shrinkage could affect cell density estimates. Although we project cell densities onto a normalized volumetric column, and although we have quantified the shrinkage of the sections, the cell density estimates might somewhat differ using alternative reconstruction methods. Another potential error could be introduced by cutting cells located at slice borders – these cells will appear in both slices, resulting in an overestimation of the cell count. We corrected for this overestimation by including only those cells within a given radius along the z-direction (which is orthogonal to the cutting plane) and no smaller than half of the average radius along x- and y-direction. This ensured that the overwhelming majority of the cells were not counted twice, as confirmed by the human observer quantifications.

### Comparison with past cell counts

In our data, the average neuronal density, as identified by NeuN staining, across all layers of the mouse barrel cortex is 1.66× 10^5^ per mm^3^, before correcting for tissue shrinkage. Assuming that each slice in our sample was cut precisely as a 50 μm section, after immunostaining the average optical thickness of slices was reduced to 32.5 μm, indicating a 34.8% shrinkage in z-direction. The shrinkage along x-y plane was generally much smaller in our protocol: imaged cells with a voxel size of 0.73-by-0.73-by-0.45 or 1.46-by-1.46-by-0.9 μm showed similar pixel radius along x-, y- and z- axes (data not shown). If we assume that the real neurons have a similar radius along the 3 axes, the data suggests a shrinkage factor of ~2.3% along x- and y- axes. After correcting for the estimated average shrinkage factors, the average neuronal density became 1.03×10^5^ per mm^3^, in agreement with the previous observations made in the C57B6 mouse (i.e. 0.6×10^5^-1.6×10^5^ per mm^3^, (Hodge et al., 2005; Irintchev et al., 2005; Lyck et al., 2007; Ma et al., 1999; Tsai et al., 2009)).

### Comparison with other simulated networks

Network models help explain network dynamics and information processing on many levels. Therefore, they exist at many different scales of complexity. On one extreme, simplified network models investigate how a single or a few aspects of the network (connectivity) properties affect network behavior. For instance, randomly connected balanced networks use integrate-and-fire neuron models (Brunel, 2000), binary neuron models (van Vreeswijk and Sompolinsky, 1998, 1996), or rate neuron models (Sompolinsky et al., 1988) to investigate the effects of synaptic sparseness, connectivity strength and the balance between excitation and inhibition on network dynamics. Similarly, like discussed in the introduction, feed-forward networks like the perceptron (Rosenblatt, 1958) can explain the increasing abstraction of receptive fields in sensory perception using similar simplified neuron models(Seung and Yuste, 2012) and randomly connected symmetric networks (Hopfield, 1982) can explain associative memory. Finally, the dynamics of small-world networks (Watts and Strogatz, 1998) have several special properties such as rapid (near-critical) synchronization, low wiring costs and a balance between locally specialized and large-scale distributed information processing (Bassett and Bullmore, 2006; Stam and Reijneveld, 2007).

Although simplified networks are often very powerful in providing (analytical) explanations about the influence of connectivity on network behavior, they are biologically not very realistic. A middle ground can be found in biologically-inspired networks that use the intrinsic connectivity schemes found in the brain. These model networks often make specific predictions about the effects of network properties on dynamics, although analytical solutions are mostly not feasible (see for instance (Rubin and Terman, 2004; Tort et al., 2007; Wendling et al., 2002)(Tort et al., 2007), (Rubin and Terman, 2004)).

Another intermediate level of network modeling involves fitting functional models to whole-network recordings (e.g. Generalized Linear Models (GLMs) (Paninski, 2004; Pillow et al., 2008; Truccolo et al., 2005), Generalized Integrate-and-Fire models (GIF models) (Gerstner and Kistler, 2002; Jolivet et al., 2004)). With these types of models, the spiking behavior and functional connectivity of entire networks can be fitted to network recordings. The results from such an analysis can be difficult to link to biophysical properties of the neurons and networks, but it is a very successful method for describing the functional connectivity of for instance the macaque, salamander, cat and rabbit retina (Denk and Detwiler, 1999; Doi et al., 2012; Keat et al., 2001; Li et al., 2015; Marre et al., 2012; Pillow et al., 2008; Reich et al., 1998) (for a review see (Field and Chichilnisky, 2007)) and C. elegans (Kato et al., 2015).

Finally, on the other extreme, are biologically reconstructed networks, like the one we present here. For some systems, complete or partial wiring diagrams have been published (C. elegans (Varshney and Beth L. Chen, 2011), mouse retina (Helmstaedter et al., 2013)), that can be used to construct such models. A notable example is the crustacean stomatogastric ganglion system, that has been extensively studied and simulated, leading to variable invaluable insights into neural network functioning in general (Marder and Goaillard, 2006; Prinz et al., 2004). These networks are biologically realistic, but because of their complexity, it is more difficult to analyze the influence of specific network properties on network dynamics and function. Moreover, one concern is that with the current methods, it is still impossible to measure all relevant parameters (molecular cell-type, electrophysiological cell-type, cell location, structural connectivity, functional connectivity) in a single sample. Therefore, every biologically reconstructed network so far is a combination of properties from different individuals and even animals. Whether such a synthesized model is a good approximation of the actual functional neural network remains to be seen (Edelman and Gally, 2001; Marder and Taylor, 2011). Moreover, all current reconstructed networks are limited in their scope: right now it is not feasible to reconstruct and model the whole brain. For the barrel cortex presented here, that means that motor and top-down input are missing, which results in reduced neural activity *in silico* than observed experimentally (compare Figure 8 and 9 to (Peron et al., 2015)) especially during hyperpolarized membrane potentials. Despite these limitations, biologically reconstructed network models are very important as a testing ground for hypotheses based on more simplified networks, or to assess biological parameters that are difficult or impossible to measure experimentally, such as the effects of threshold adaptation (Huang et al., 2016; Zeldenrust et al., 2020) or the effects of different coding schemes (Huang et al., 2020). In Supplemental Table 1, we have summarized the properties of several biologically reconstructed networks that have been published. Note that until now, many of these reconstructed networks have to be run on a cluster of computers or on a supercomputer, because a simple desktop computer simply lacked the computational power to run a biologically reconstructed network and/or did not make the code available (Tomsett et al., 2015) being an exception). We used simplified neuron models instead of reconstructed multi-compartmental models, increasing the computational efficiency, but possibly missing effects due to the morphology, such as certain forms of bursting (Zeldenrust et al., 2018), dendritic computation (Chu et al., 2020) or axon-initial segment effects (Kole and Brette, 2018). Finally, like the recent model by Markram et al. (Markram et al., 2015), we used no parameter tuning to construct this model, other than making the different cell-types of the Izhikevich-model and controlling the cell-type specific connection probabilities. All this makes the model very accessible for quickly testing fundamental hypotheses systematically (Huang et al., 2020, 2016).

## Materials and Methods

### Experimental procedures

#### Tissue preparation and immunochemistry

The slices from the barrel cortex were described before (Kole et al., 2020; Kole and Celikel, 2019) with minor modifications. In short, juvenile mice from either sex were perfused using 4% paraformaldehyde before tangential sections were prepared. To ensure that cortical layers were orthogonal to the slicing plane the cortex was removed from the subcortical areas and medio-lateral and rostro-caudal borders trimmed. The remaining neocortex included the entire barrel cortex and was immobilized between two glass slides using four 1.2 mm metal spacers. The rest of the histological process, including post-fixation and sucrose treatment, was performed while the neocortex was flattened. All care was given to ensure that the tissue is as flat as possible at the time of placement onto the sliding horizontal microtome. 50-micron sections were cut and processed using standard immunohistochemical protocols. The following antibodies were used: anti-NeuN (Millipore, Chicken), anti-GAD67 (Boehringer Mannheim, Mouse), anti-GABA (Sigma, Rabbit), anti-Parvalnumin (PV, Swant Antibodies, Goat), anti-Somatostatin (SST, Millipore, Rat), anti-Calretinin (CR, Swant Antibodies, Goat), anti-vasointestinal peptide (VIP, Millipore, Rabbit) at concentrations suggested by the provider.

The imaging was performed using a Leica Confocal microscope (LCS SP2) with a 20X objective (NA 0.8). Each section sequentially cutting across layers was individually scanned with 512×512 pixel resolution; the signal in each pixel was average after 4 scans and before it was stored. The alignment of each section was performed automatically using a fast Fourier transform based image registration method (Guizar-Sicairos et al., 2008)

#### Automated cell counting

All image analysis was done using a custom-written running toolbox in Matlab 2012b with an Image Processing Toolbox add-on (Mathworks).

#### Nucleus-staining channels (NeuN, Parvalbumin and Calretinin)

Most fluorescence imaging methods, including confocal microscopy, have several shortcomings that make the automated cell identification a challenging task: First, the background intensity of images is often uneven due to light scattering and tissue auto-fluorescence. Shading and bleaching of fluorophores further add to this problem when acquiring multiple confocal images at the same location. Second, intensity variation within a single cell might cause over-segmentation of the cell. Third, the intensity of different neuron populations turn out to be very different because they absorb fluorescent dye unevenly. Specifically, GAD67+ and SST+ neurons usually have a weakly stained nucleus as visualized by anti-NeuN antibody, making non-linear gain modulation necessary in a cell-type specific manner. To overcome these problems and maximize the hit and correct rejection rate over miss and false positives (i.e. (H+CR)/(M+FP)), we have developed the following pipeline:

##### Pre-processing

The goal of pre-processing is to obtain relatively consistent images from original fluorescent images with varying quality to pass to the cell count algorithm, so the same algorithm can process a large variety of images and still get consistent results. Depending on the nature of the individual channel, i.e. which antibody was used, different pre-processing steps were employed.

##### Median filtering

A median filter with 3×3×3 pixel neighborhood is applied to fluorescent image stacks to smooth intensity distribution within each image stack in 3D. This operation removes local high-frequency intensity variations (Supplemental Figure 1b).

##### Vignetting correction

Vignetting is the phenomenon of intensity attenuation away from the image center. We use a single-image based vignetting correction method (Zheng et al., 2009) to correct for the intensity attenuation (Supplemental Figure 1c). The algorithm extracts vignetting information using segmentation techniques, which separate the vignetting effect from other sources of intensity variations such as texture. The resulting image is the foreground, i.e. the cellular processes, on a homogenous background.

##### Background subtraction

Background can result from non-specific binding of antibodies or auto-fluorescence of the tissue. To reduce the background noise, local minima in each original grayscale image are filled by morphological filling, and background is estimated by morphological opening with 15 pixel radius disk-shaped structuring element. The radius value is chosen to be comparable to the largest object size so the potential object pixels are not affected. The estimated background is then subtracted from the original image to enhance signal-to-noise ratio, SNR (Supplemental Figure 1d).

##### Contrast-limited adaptive histogram equalization (CLAHE)

CLAHE (Heckbert, 1994) enhances local contrast within individual images by remapping intensity value of each pixel using a transformation function derived from its neighbourhood. While increasing local contrast and amplifying weakly stained cells, it also reduces global intensity difference, which partially corrects for the uneven illumination that individual fluorescent images often suffer from (Supplemental Figure 1e). CLAHE is applied as an 8×8 tiles division for each image. Images from channels with very low number of positive staining with high SNR (e.g. Calretinin staining channel) are not processed with CLAHE.

###### Image segmentation to identify cell nucleus

Black-and-white image transform is applied to grayscale images to separate foreground, i.e regions presumably contain nuclei, from background. In the ideal conditions, if all the objects were stained evenly during immunochemistry, the image pixels’ intensity value will be distributed as two well-separated Gaussian distributions. However, objects are usually not evenly stained; specifically, GAD67+ and SST+ neurons usually have weak NeuN staining. As a result, the intensity distribution for object pixels is very broad and cannot be described by a single Gaussian distribution. To reliably identify foreground pixels we calculated threshold values using 2-level Otsu’s method (Otsu, 1979), which separates the pixels into 3 groups. The group with the lowest intensity reliably captures the background pixels, and the other 2 groups are set to the foreground. This transformation is directly applied to 3D image stack to obtain 3D foreground (Supplemental Figure 1f).

##### Marker-based watershed segmentation

B&W transform identified regions contains cell nucleus, albeit non-specifically, and it does not identify the location and shape of each individual nucleus stained, thus image segmentation is needed to identify individual nuclei. Watershed method (Meyer, 1994) is an efficient way of segmenting grayscale images, i.e. foreground part of image obtained by B&W transformation based on gradient, and has the advantage of operating on local image gradient instead of global gradient. However, direct application of watershed methods usually results in over-segmentation of nuclei due to local intensity variation within individual nuclei. To overcome this problem, marker-based watershed algorithm is employed, in which markers serving as starting ‘basin’ for each object are first placed on an image to be segmented, and watershed algorithm is then applied to produce one segment (or object) on each marker.

We computed the markers by applying regional maxima transform on foreground grey-scale images. To ensure at most one marker is placed in each nucleus, first the grey-scale image need to be smoothed to eliminate local intensity variation. This is realized by applying morphological opening-by-reconstruction operation (Vincent, 1993) with 5 pixels radius on foreground grayscale image, which removes small blemishes in each individual nucleus and ensures regional maxima transform can find foreground markers accurately.

After identifying markers watershed algorithm is applied (Supplemental Figure 1g). To ensure accurate detection of cell boundaries, the B&W foreground needs to enclose the entire cell object. This image dilation is applied to the B&W foreground to enlarge it by 1 pixel in radius before application of watershed segmentation algorithm. Finally, objects with size smaller than 400 pixels in total are removed by morphological opening.

##### Corrections for clusters of connected neurons

Clusters of closely located neurons are not always successfully separated without further image processing; especially when closely located neurons all have similar intensity distribution. In such cases application of intensity-based watershed algorithms result in identification of one object instead of many real neurons (Supplemental Figure 1b). Furthermore, our strategy for watershed segmentation to augment regional intensity similarity to make sure that nuclei are over-segmented actually increases the chance of under-segmentation during clustering. To correct for this under-segmentation we employed a five-step approach:

a. The volume (total number of pixels) of all identified objects is calculated, and objects with a volume larger than mean+std of the population are labeled as “potential clusters”.
b. For each object in the potential cluster list, the original grayscale image is retrieved. Then, from all the pixels contained in the object, 50% pixels with lower intensity values are removed, generating a new B&W object with a smaller size. Because usually, those low-intensity pixels are from the periphery region of each individual neuron, the new B&W object has better separation between different neurons.
c. Euclidean distance-based 3-D regional maximum transform is then applied to the new, smaller B&W 3-D candidate object, in which the distance from each pixel belongs to the object to the border of the object, is calculated. Assuming neurons have Ellipsoid-like shape, the peak (largest distance from borders) of this transform will likely be the center of neurons, even if they are connected. The regional maximum transform is then applied to locate those peaks in the Euclidean distance space. Before the regional maximum transform is applied, the target image is smoothed by morphological opening-by-reconstruction operation with 1-pixel radius to remove small local variations.
d. If more than one center is found (in c) watershed method is applied to the distance transform of the original B&W object, using the identified centers as markers. If only one center is found then the cluster is judged as a single neuron and removed from the list. Again, the distance metric is smoothed by a morphological opening-by-reconstruction operation before the watershed algorithm is applied.
e. Steps a-d is repeated until the “potential cluster list” is empty (Supplemental Figure 1h).

##### Morphological filtering

Neurons have a certain shape and volume. Based on this statistical information clustered objects can be filtered to remove small artifacts. This is necessary because of the low threshold value used for the foreground generation. To remove the artifacts from neurons we first performed a morphological opening with a structure whose size is 1/3 of the size of each object’s bounding box. The bounding box is calculated in 3-D hence it is the smallest cube that contains the object. This operation breaks down irregular shapes but keeps relatively regular shapes (sphere, ellipsoid, cuboid) intact. Then, both pixel size (volume) and mean intensity of the objects are fitted with a Gaussian mixture model, and the group with the smallest pixel size and lowest mean intensity is judged as an artifact and is removed. (Supplemental Figure 1f).

##### Combining information from different soma-staining channels

Cells identified from each channel are added together to give cumulative soma counts across all antibody channels. Overlapped objects are judged to be different cells if:

a. Overlapping is smaller than 30% of any object volume constituting the cluster
b. after subtraction the new object preserves the ellipsoid shape

#### Cytosol-staining channels (GAD67 and Somatostatin)

Identification of the cells in cytosol-staining channels utilizes reference information gathered from the soma-staining channels, hence segmentation of cytosolic signals requires at least one nuclear channel staining.

Early stages of the image processing for the cytosolic signal localization was identical to that of soma-staining channels except CLAHE step. Subsequently, cell objects were imported from combined soma-staining channels information (Supplemental Figure 2c).

For each cell object, two additional pixels were added to the diameter of the object (Supplemental Figure 2d). This enlarged cell object is used as a mask to detect positive staining in the cytosol-staining channel (Supplemental Figure 2f). Positive staining was defined as connected pixels with a volume at least 10% of the object and that they have significantly higher intensity compared to the pixels within 2.5 times of the associated cell (Supplemental Figure 2g). Finally, the percentage of positive staining was obtained and used to identify GAD67 or Somatostatin positive cells.

##### Performance comparison between computer and the human observer

Three human observers independently counted a number of 3-D images stacks from different antibody staining, using Vaa3D software (Peng et al., 2010). Three identical copies of each image stack were placed in the manual counting dataset in random order; the human observers subsequently confirmed that they did not notice the duplicates in the data set they had analyzed. The automated counting result was compared with the average human counting result, and the summary of the difference is shown in Supplemental Table 5.

#### Generating an average barrel column

After performing automatic cell counting on individual slices across different cortical depths, we calculated average cell density for different types of cells identified by distinct antibody channels at a given cortical depth as indicated by slice number. Tissue shrinkage was not corrected but the average column size was empirically determined. To account for the differences in cortical thickness across different animals, we then binned the density data from each individual animal into 20 bins, which were subsequently averaged to obtain the average cell density distribution across cortical depth. The layer borders *z_lim_* between different cortical layers (L1-L2/3, L2/3-L4, L4-L5, L5-L6) were determined as described previously (Meyer et al., 2010), by first fitting a Gaussian function

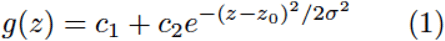

to the NeuN+ cell density profile along with cortical depth with manually set *c*_1_, *c*_2_ and *z*_0_, and then the respective *z_lim_* was calculated as

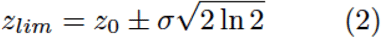

L5A-L5B border was determined by manual inspection on NeuN+ cell density. We then calculated the size of an average barrel in C-E rows, 1-3 columns by manually labeling corresponding barrels in anti-GAD67 staining (Supplemental Figure 3). The number of different types of cells in an average barrel from C-E rows, 1-3 columns was then calculated by the size as well as the corresponding cell density.

### Network setup

#### Neuronal Model

We used the Izhikevich quadratic model neuron (Izhikevich, 2004, 2003) in this study:

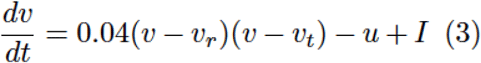

where *v, v_r_*, and *v_t_* are the membrane potential, resting membrane potential without stimulus, and the spike threshold of the neuron, respectively and *I* is the synaptic current the neuron received (see below). The dynamics of the recovery variable u are determined by:

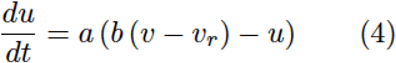

Parameters *a, b, c, d* together determine the firing pattern of the model neuron (see Supplemental Table 4). The model has the following reset condition:

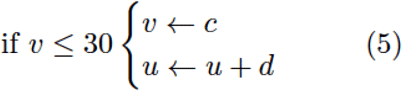

Parameters *a, b* and *c* were taken from (Izhikevich, 2003); parameter *d* was adapted to match firing rates observed in the literature (see 4.2.2). For the simulations, a first-order Euler method with a step size of 0.1 ms was used.

#### Neural Network Model

##### Neural Distributions

The mouse barrel cortex L4-L2/3 network is modeled based on the distribution of different classes of neurons in an average barrel reconstructed by immunochemical labeling and confocal microscopy (see above). 13 different types of cortical neurons are included in the model (Markram et al., 2004; Oberlaender et al., 2012; Thomson and Lamy, 2007). In L2/3 there are 9 types of neurons, 2 excitatory: L2 pyramidal neurons and L3 pyramidal neurons (Brecht et al., 2003; Feldmeyer et al., 2006); 7 inhibitory: PV+ fast-spiking neurons (Holmgren et al., 2003; Packer and Yuste, 2011), PV+ bursting neurons (Blatow et al., 2003), SST+ Martinotti neurons (Fino and Yuste, 2011; Kapfer et al., 2007; Wang et al., 2004), Neurogliaform cells (Tamás et al., 2003; Wozny and Williams, 2011), CR+ bipolar neurons (Caputi et al., 2009; Xu et al., 2006), CR+/VIP+ multipolar neurons (Caputi et al., 2009) and VIP+/CR-neurons (Porter et al., 1998). In L4 there are 4 types of neurons, 2 excitatory: L4 spiny stellate neurons and L4 star pyramidal neurons (Egger et al., 2008; Staiger et al., 2004); 2 inhibitory: PV+ fast-spiking neurons and PV- low-threshold spiking neurons (Beierlein et al., 2003; Koelbl et al., 2015; Sun et al., 2006). The distribution of excitatory, PV+, CR+, and SST+ neurons are taken from the anatomical reconstructions; for other cell types, we assigned corresponding number of different neurons in each cluster based on the previous studies (Kawaguchi and Kubota, 1997; Uematsu et al., 2008). These neurons were distributed in a 640-by-300-by-300 μm region (L4, 210-by-300-by-300; L2/3, 430-by-300-by-300). Note that we scaled the size of the network to match the average dimension of a rat column (Feldmeyer et al., 2006), due to the fact that most of the axonal and dendritic projection patterns were measured in the rat.

##### Connectivity

Connectivity is determined using axonal and dendritic projection patterns (Egger et al., 2008; Feldmeyer et al., 2006, 2002; Helmstaedter et al., 2008; Lübke et al., 2003) which are approximated by 3-D Gaussian functions, with the assumption that the probability that two neurons are connected is proportional to the degree of axonal-dendritic overlap between these two neurons (i.e Peter’s rule, (White, 1979)). For each pre-synaptic *i* and post-synaptic neuron *j*, we calculate the axonal-dendritic overlapping index *I_i,j_*, which is the sum of the product of presynaptic axonal distribution and postsynaptic dendritic distribution *D_j_*:

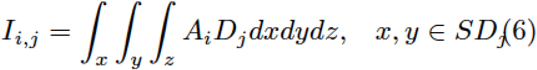

where *SD_j_* is the 3-D space that contains 99.9% of *D_j_*. We then convert *I_ij_* into connection probability *P_i,j_* between neuron *i* and *j*, by choosing a constant k for each unique pre- and post-synaptic cell type pair so that the average connection probability within experimentally measured inter-soma distances (usually 100 μm) matches the empirically measured values between these two types of cells (Supplemental Table 3):

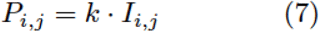

Finally, a binary connectivity matrix was randomly generated using the pairwise connection probabilities *P_i,j_*, in which connected pairs are labeled as 1.

##### Synapses

Synaptic currents in this network are modeled by a double-exponential function. Parameters of those functions are adjusted to match experimentally measured PSPs (peak amplitude, rise time, half-width, failure rate, coefficient of variation and pair-pulse ratio) in the barrel cortex *in vitro* (Supplemental Table 3; see (Thomson and Lamy, 2007) for an extensive review). The onset latency is calculated from the distance between cell pairs; the conduction velocity of the action potential was set to 190μm/ms (Feldmeyer et al., 2002). The short-term synaptic dynamics (pair-pulse depression/facilitation) is modeled as a scalar multiplier to actual synaptic weight, which follows a single exponential dynamic (Izhikevich and Edelman, 2008):

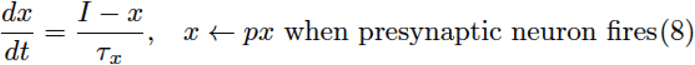

*τ_x_* was set to 150ms for excitatory synapses and depression inhibitory synapses (*p*<1), and 100ms for facilitating inhibitory synapses (*p*>1). Differences in the activation state of cortex are included in the model by setting the common initial voltage and the equilibrium potential *v_r_* of all cells, thus accounting for potential up - and down-states as well as an intermediate state.

##### Thalamic inputs into the barrel cortex *in silico*

To the best of our knowledge, there is not any published quantitative work on the cellular organization of the mouse thalamic nuclei. In the rat, each barreloid in thalamic VPM nuclei has ~1/18 number of neurons compared to the corresponding L4 barrel (Meyer et al., 2013). Given that in our average barrel column L4 contains ~1600 neurons, we assigned between 100 and 200 thalamic neurons to each barreloid in VPM. The thalamic-cortical connectivity is calculated using the same method as cortical-cortical connectivity discussed above, using published thalamic axon projection patterns (Furuta et al., 2011; Oberlaender et al., 2012). The POM pathway was not modeled.

Each of the thalamic neurons is modelled as a ‘filter and fire’ neuron (Chichilnisky, 2001; Keat et al., 2001; Pillow et al., 2008; Truccolo et al., 2005), where each of the thalamic neurons responds to either whisker angle (filters and activation functions randomly chosen based on a parametrization of the filters from (Petersen et al., 2008)), curvature, or a combination of both. The center barreloid was considered to be the principal barreloid for the spared whisker, whereas the other two were considered secondary barreloids, which meant that they received the stimuli reduced (30% of original amplitude) and delayed (2.5 ms) (Brecht et al., 2003; Brecht and Sakmann, 2002). The thalamic spike trains served as input to the cortical model, which similarly consisted of three cortical columns, corresponding to the three thalamic barreloids. An example of how to run these simulations can be found on Github: https://github.com/DepartmentofNeurophysiology/Cortical-representation-of-touch-in-silico.

Thalamic stimulation in the model based on population PSTHs (Figures 2–7) was collected extracellularly in anesthetized animals in *vivo* (Aguilar, 2005). The PSTHs only specified the population firing rate in the thalamic cells; to generate individual neuron response in different trials we assume that thalamic neurons fire independent Poisson spike trains in each trial, constrained by the PSTHs.

#### Spike-timing dependent plasticity

A network of 3 barrel columns, representing canonical C,D,E rows, was constructed to simulate spike-timing-dependent plasticity in the barrel cortex following a single (D-row) whisker deprivation. Each column was randomly generated using distributions of 13 different types of neurons, and connectivity was calculated using the same method discussed above. The middle column was whisker-deprived, which received surround whisker evoked thalamic input; the two lateral columns were whisker-spared and received principal whisker evoked thalamic input (Aguilar, 2005). The STDP rule for L4-L2/3 excitatory connections was as follows (Celikel et al., 2004):

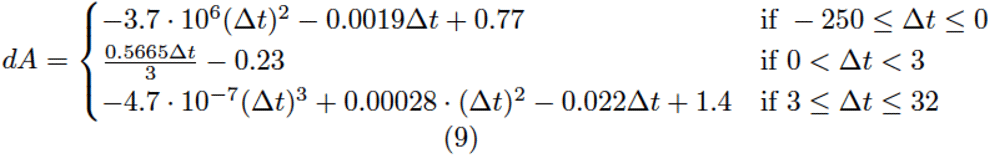

*Δt* was the timing difference (in ms) between the time at which presynaptic spike arrives at postsynaptic neuron (i.e. presynaptic neuron spike time plus synaptic delay) and the time at which the postsynaptic neuron spikes ms. The constants were directly taken from the literature, in which the values were obtained by least-square fits to the experimental data. For L2/3-L2/3 excitatory connections, the rule was as follows (Banerjee et al., 2014):

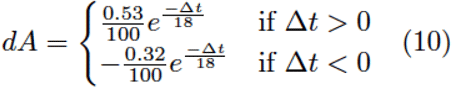

The synaptic weight change was additive for potentiation and multiplicative for depression; repeating the simulations with an additive rule for potentiation and depression did not change the results and are not shown herein. Plasticity rules for excitatory-inhibitory and inhibitory connections are less commonly studied. Inclusion of the empirically identified learning curves (Haas et al., 2006; Lu et al., 2007) did not qualitatively alter the results and are not included herein.

#### Simulated freely whisking experiment

In the simulations of a freely whisking experiment, the network (Figure 8: 3 barrels, Figure 9: 1 barrel) was presented with the whisker angle and curvature recorded from a freely moving rat (animal an171923, session 2012_06_04) in a pole localization task (data from (Peron et al., 2015) made available as ‘ssc-2’ on CRCNS.org).

NB Direct whisker modulation by motor cortex (Crochet et al., 2011) can be optionally included in the model, but was not used for our current simulations. However, it is present in the online code as option.

## Acknowledgements

This work was supported by grants from the European Commission (Horizon2020, nr. 660328), European Regional Development Fund (MIND, nr. 122035) and the Netherlands Organisation for Scientific Research (NWO-ALW Open Competition, nr. 824.14.022) to TC and by the Netherlands Organisation for Scientific Research (NWO Veni Research Grant, nr. 863.150.25) to FZ.

