## Supplemental Figures and Tables for "Cortical representation of touch *in silico*"

### 1. Supplemental Figures

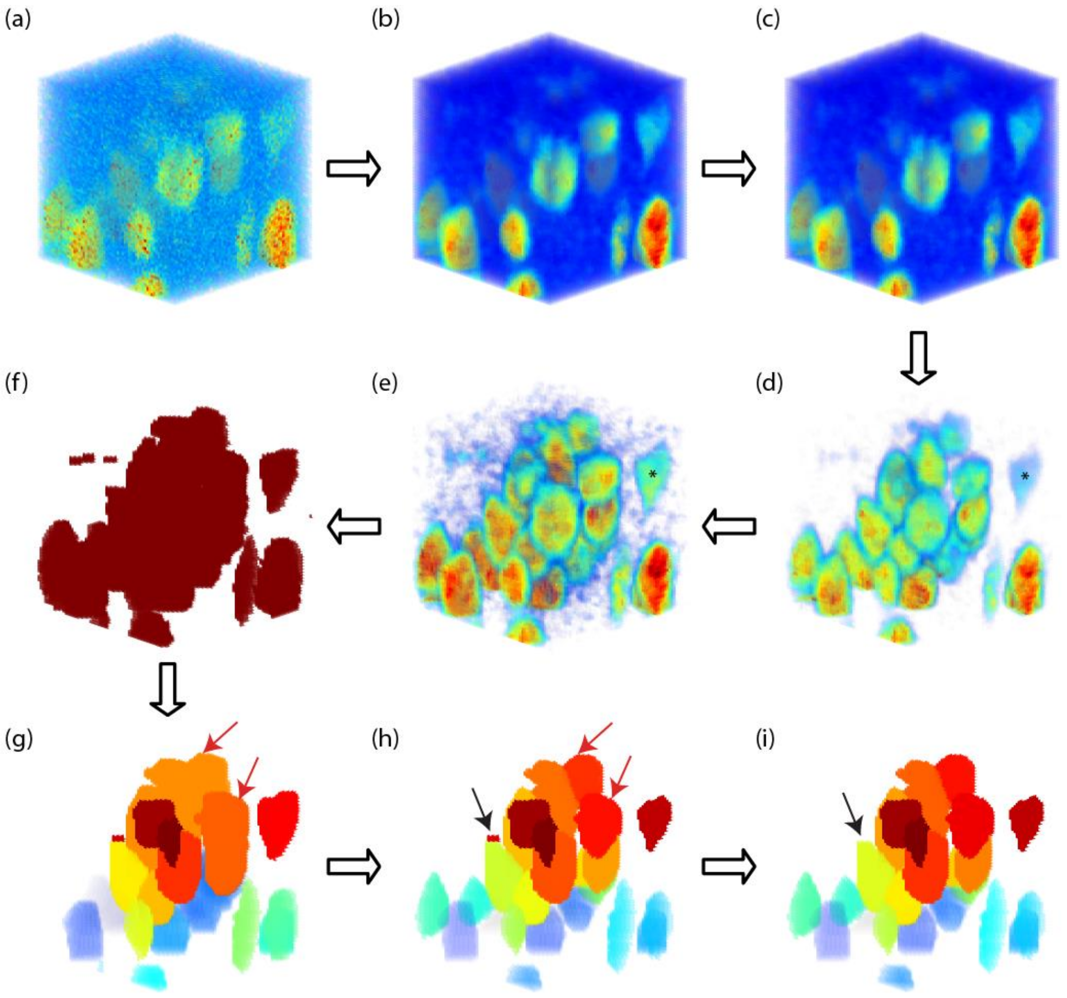


**Supplemental Figure 1.**

**Automatic cell counting for soma-staining channels.** The example is an anti-NeuN+ staining volume. (a) Original 3D image stack obtained from Z-scan confocal imaging. (b) Median filtering with a 3-by-3-by-3 pixel neighborhood to reduce local intensity variation in the image. (c) Vignetting correction using single-image based method. (d) Background subtraction to reduce the background intensity level. (e) Contrast-limited adaptive histogram equalization (CLAHE) to enhance local contrast. Notice the contrast enhancement effect on the weakly stained cell marked with * in (d) and (e). (f) Black-and-white (B&W) transform to separate foreground objects from the background. (g) Marker-base watershed segmentation on the BW image. Identified objects are labelled with different colors. Notice the cell clusters pointed with red arrows, which are under-segmented by the watershed algorithm. (h) Cluster-separation using a Euclidean distance transform. The under-segmented clusters in (g) are further segmented, as the red arrows pointed out in (g). (i) Morphological filtering on the identified objects. Small artefacts, such as the one indicated by black arrow in (h), are removed.

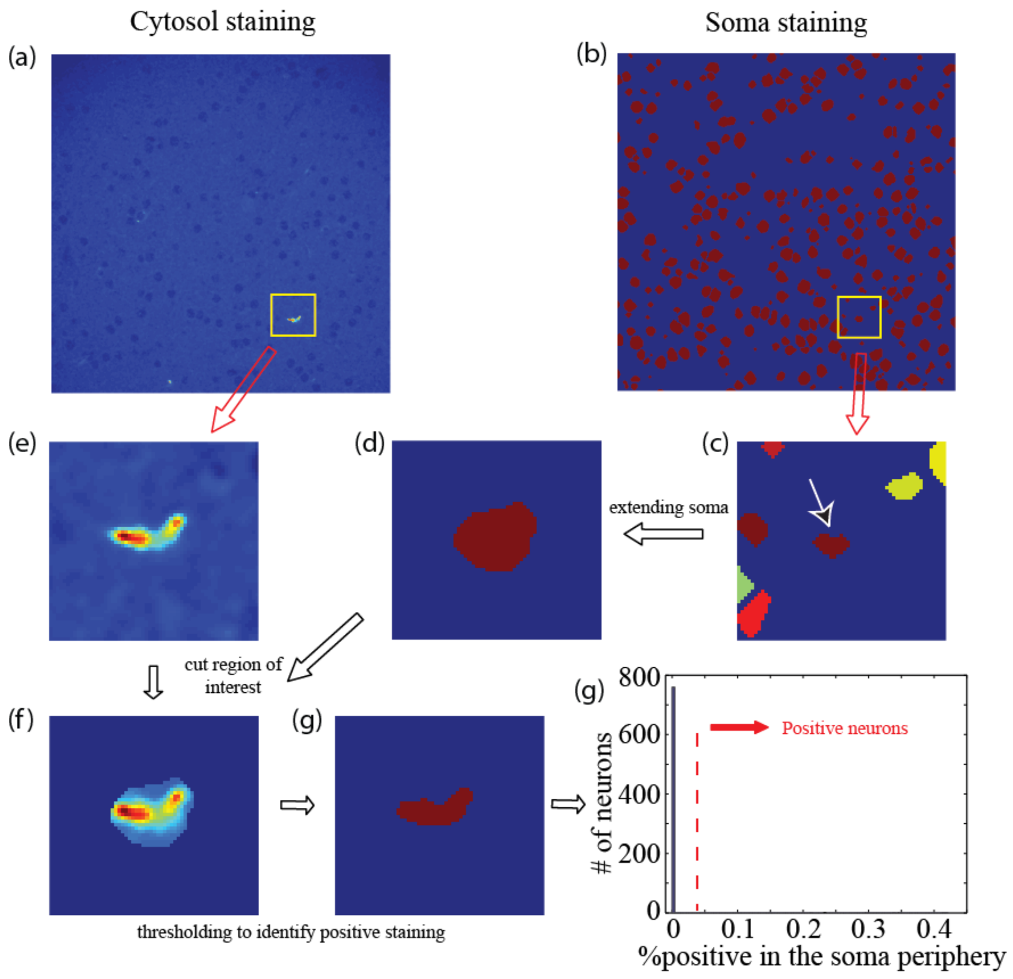


**Supplemental Figure 2.**

**Automatic cell counting for cytosol-staining channels.** The example shown is a Somatostatin (SST) staining immunofluorescent image. (a, b) Images from SST-channel and identified cell objects from soma-staining channels, at the same position in the image stack. (c) The cell object under analysis, as indicated by the red arrow, with region of interest marked by yellow rectangle in (a) and (b). (d) The cell object under analysis is enlarged by 2-pixels in all directions; other objects are ignored. (e) Corresponding region in the SST channels. (f) The object in (d) is used as mask image to obtain the region of interest in the SST channel. (g) Positive staining is detected as connected pixels with at least 10% volume of object in (d), whose intensity value is two times the standard deviation higher than average pixel intensity in the local image region as shown in (c). (h) Histogram of percentage of positive staining in the SST channel for all the cell objects identified in (b). Cells with a positive percentage higher than 9% are labelled as SST-positive cells.

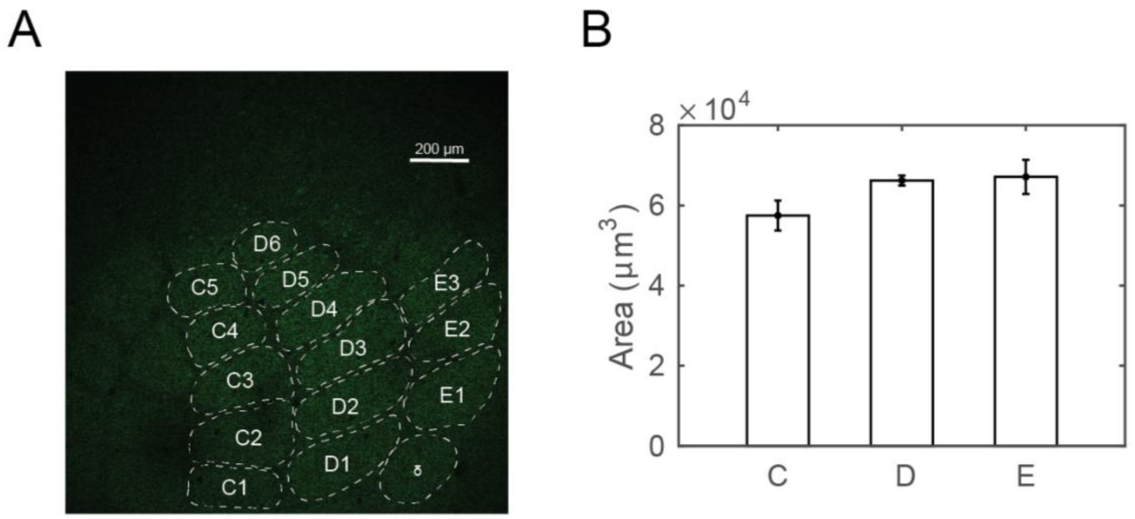


**Supplemental Figure 3.**

**Average volume of a cortical column.** (A) A sample image stained with anti-GAD67 antibody overlaid with manually determined barrel borders. (B) Average area of the Layer 4 across barrel columns C, D, and E.

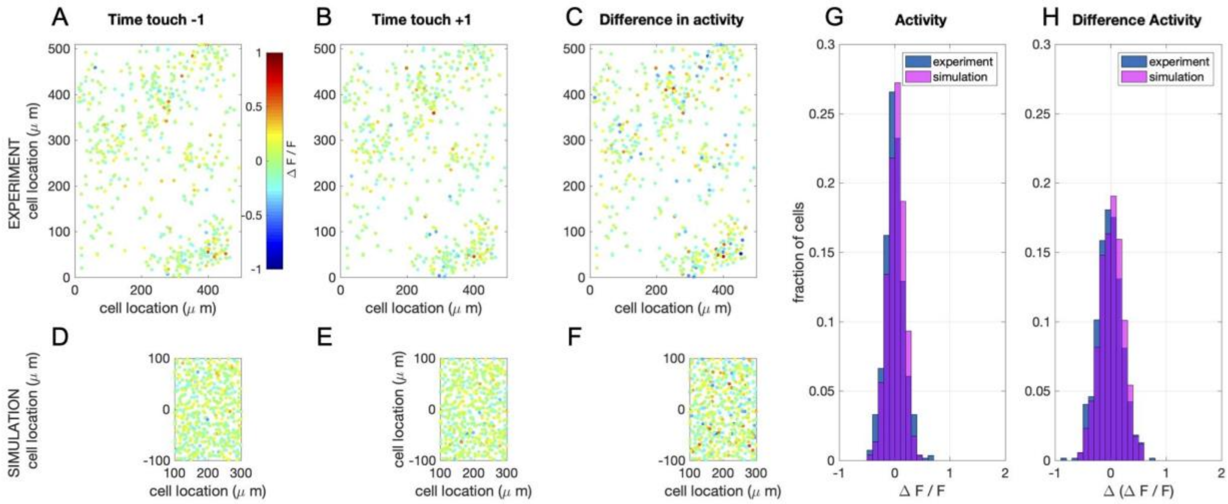


**Supplemental Figure 4.**

**Simulation of calcium imaging experiment in L4 (see main text Figure 9). (A)** Recorded network response 1 time frame before touch (sampling frequency: 7 Hz; recorded volume: 8). **(B)** Recorded network response 1 time frame after touch **(C)** Difference in network response between before and after touch. **(D-F)** Same as in A-C, but now for simulations (full simulation: single barrel including L23 and L4 (shown here)). Note that a recorded volume is larger than a single barrel. The frames are scaled accordingly. **(G)** Comparison of the distribution of activity of 1 time frame after touch between the recorded and the simulated network. **(H)** Comparison of the distribution of the difference in activity between 1 time frame before and after touch between the recorded and the simulated network.

### 2. Supplemental Tables

| Citation | Brain regions | Model neurons | Number of neurons in the model | Synaptic plasticity | | Comp. platform | Open source toolbox |
|---|---|---|---|---|---|---|---|
| | | | | Short term | Long term | | |
| Traub et al. 2005 | Rat thalamus & cortex | MC conductance | 3560 | No | No | Cluster | No |
| Markram et al. 2006 | Cortical column | MC conductance | 10K | Yes | No | Blue Brain infrastructure, supercomputer | No |
| Izhikevich & Edelman 2008 | Mammalian brainstem, thalamus, cortex | Izhikevich neuron - MC | 1M | Yes | STDP | Cluster | No |
| Ananthanarayanan et al. 2009 | Cat visual thalamus and cortex | Izhikevich neuron - Point neuron | >1M | No | STDP | Super computer | No |
| Zhu et al. 2009 | Macaque V1, L4 | Leaky integrate and fire | 16384 | No | No | Unknown | No |
| Phoka et al. 2012 | Rodent barrel cortex L2-4 | Izhikevich neuron - MC | 3717 | No | STDP | NEST | No |
| Reimann et al. 2013 | Rodent neocortical column | MC conductance | hundreds | No | No | NEURON, Blue Brain infrastructure, supercomputer | No |
| Potjans & Diesmann, 2014 | Cortical microcircuit | Leaky integrate-and-fire | 80K | No | No | NEST, cluster | Yes (NEST website) |
| Sharp et al. | Rodent | Leaky | 50K | No | No | Cluster | No |

| | | | | | | | |
|---|---|---|---|---|---|---|---|
| 2014 | barrel cortex | integrate and fire | | | | (SpiNNaker) | |
| Markram et al. 2015 | Rat thalamus and hindlimb S1 | MC conductance | 31K | Yes | Yes* | Super computer | No |
| Tomsett et al. 2015 | Macaque neocortical network | Reduced MC conductance | >100K | No | No | Matlab, desktop computer | Yes (ModelDB) |
| Chariker et al. 2016 | Macaque V1 | Leaky integrate-and-fire | 4K | No | No | Unknown | No |
| Sudhakar et al. 2017 | Cerebellar granular layer | MC conductance | 800K | No | No | NEURON, cluster | Yes (ModelDB) |
| Smith et al. 2018a, Schmidt et a. 2018b, Schuecker et al. 2017 | Macaque visual cortex, long range connectivity | Leaky integrate and fire | ~300K | No | No | NEST, Juqueen supercomputer | Yes (https://inm-6.github.io/multi-area-model/) |
| Allen Brain Arkhipov et al. 2018, Billeh et al. 2019 | Mouse thalamus, L4 visual cortex | Hodgkin-Huxley style conductance and leaky integrate-and-fire | 230K | No | No | Cluster (HH) / desktop (IF) | Yes (brain-map.org) |
| Bernardi et al. 2020 | Rodent barrel cortex | Leaky integrate and fire | 2600 | Yes | No | Unknown | No |
| Huang et al (Current article) | Rodent thalamus L2-4 barrel cortex | Modified Izhikevich neuron - Point neuron | 4200 (depends on number of columns created) | Yes | STDP | Matlab, desktop computer | Yes (on GitHub) |

**Supplemental Table 1.**

**Summary of bioinspired cortical network models.** MC: Multi compartmental. S1: Primary somatosensory cortex. * The type of long-term synaptic plasticity model is unknown.

| | Layer 2/3 | Layer 4 | Layer 5a | Layer 5b | Layer 6 |
|---|---|---|---|---|---|
| **NeuN+** | 2410 ± 154 | 1557 ± 192 | 709 ± 133 | 811 ± 68 | 1226 ± 232 |
| **GAD67+** | 6.5 ± 0.7% | 5.5 ± 0.6% | 9.5 ± 1.7% | 10.2 ± 3.9% | 6.8 ± 1.7% |
| **PV+** | 5.3 ± 1.2% | 3.3 ± 0.4% | 13.0 ± 5.6% | 9.4 ± 3.1% | 5.3 ± 0.3% |
| **SST+** | 1.5 ± 0.5% | 1.5 ± 1.1% | 3.1 ± 3.4% | 1.7 ± 1.8% | 1.7 ± 1.0% |
| **CR+** | 0.9 ± 0.1% | 1.2 ± 0.2% | 0.8 ± 0.5% | 0.4 ± 0.2% | 0.6 ± 0.5% |

**Supplemental Table 2**

**Distribution of distinct cell populations in a canonical (D-row) barrel cortical column.** Values are mean ± std. The estimates are based on 3D reconstructions across five animals, except in Layer 5b (N=3) and Layer 6 (N=2). All values, except NeuN+, are in respect to the NeuN+ cell count within the corresponding layer.

| Presyn. neuron | Postsyn. neuron | Ampli-tude (mV) | Rise time (ms) | Decay time (ms) | PPR | Failure rate (%) | CV | Hit rate | Ref ** |
|---|---|---|---|---|---|---|---|---|---|
| **Thalamic projections into the L4** | | | | | | | | | |
| Thalamic | Cortical excitatory | 0.95 ± 1.10 | 1.16 ± 0.27 | 22.5 ± 27.4 | 0.76 ± 0.07 | 0.00 ± 0.01[2] | 0.23 ± 0.15[2] | 0.43[3] | A, B, C |
| | Cortical fast spiking | 1.62 ± 1.26[1] | 0.41 ± 0.15 | 6.74 ± 1.10 | 0.55 ± 0.12 | 0.00 ± 0.01[2] | 0.22 ± 0.12[2] | 0.5[3] | |
| | Cortical non-fast spiking | 0.27 ± 0.19[1] | 1.12 ± 0.48 | 22.5 ± 11.10 | 0.76 ± 0.07 | 0.00 ± 0.01[2] | 0.72 ± 0.34[2] | 0.5[3] | |
| **L4 – L4 connections** | | | | | | | | | |
| L4 excitatory | L4 excitatory | 1.1 ± 1.1 | 0.88 ± 0.26 | 12.3 ± 2.2 | 0.65 ± 0.16 | 0.11 ± 0.18 | 0.30 ± 0.19 | 0.06 | A |
| | L4 fast spiking | 2.2 ± 2.2 | 0.37 ± 0.11 | 4.9 ± 1.9 | 0.65 ± 0.16 | 0.03 ± 0.08 | 0.27 ± 0.13 | 0.43 | |
| | L4 low-threshold spiking | 0.3 ± 0.5 | 0.86 ± 0.48 | 8.9 ± 2.9 | 1.2 ± 0.3 | 0.57 ± 0.35 | 1.04 ± 0.54 | 0.57 | |
| L4 fast spiking | L4 cells | 1.1 ± 0.8 | 1.5 ± 0.7 | 24.0 ± 10.8 | 0.72 ± 0.25 | 0.03 ± 0.07 | 0.25 ± 0.11 | 0.44 | |
| L4 low-threshold spiking | L4 cells | 0.48 ± 0.45 | 2.1 ± 1.0 | 22.6 ± 13.7 | 0.99 ± 0.26 | 0.29 ± 0.26 | 0.41 ± 0.21 | 0.35 | |
| **L4 – L2/3 connections** | | | | | | | | | |
| L4 excitatory | L2/3 pyramidal | 0.7 ± 0.6 | 0.8 ± 0.3 | 12.7 ± 3.5 | 0.90 ± 0.39 | 4.9 ± 8.8 | 0.27 ± 0.13 | 0.12 | D |
| | PV+ fast-spiking cell | 0.96 ± 0.93 | 0.89 ± 0.31 | 15.0 ± 8.2 | 0.99 ± 0.66 | 20.0 ± 20.0 | 0.27 ± 0.13* | 0.2[4] | E, F |
| | PV+ bursting cell | 1.2 ± 0.2 | 0.42 ± 0.1 | 6.3 ± 2.1 | 0.84 ± 0.17 | 13.0 ± 19.8 | 0.5 ± 0.3* | | |
| | Martinotti neuron | Not connected | | | | | | | |
| | Neurogliaform cell | 0.59 ± 0.21 | 0.85 ± 0.26 | 13.0 ± 6.2 | 0.70 ± 0.18 | 17.0 ± 14.0 | 0.5 ± 0.3* | 0.2[4] | |

| | | | | | | | | | |
|---|---|---|---|---|---|---|---|---|---|
| | CR+ bipolar cell | 1.3 ± 0.9 | 1.1 ± 0.4 | 14.0 ± 4.1 | 1.00 ± 0.62 | 13.0 ± 19.8 | 0.5 ± 0.3* | | |
| | CR+ multipolar cell | 1.4 ± 1.4 | 0.79 ± 0.50 | 8.5 ± 3.1 | 0.92 ± 0.40 | 13.0 ± 19.8 | 0.5 ± 0.3* | | |
| | VIP+/CR- cell | 1.3 ± 0.9 | 1.1 ± 0.4 | 14.0 ± 4.1 | 1.00 ± 0.62 | 13.0 ± 19.8 | 0.5 ± 0.3* | | |
| L4 fast spiking | L2/3 cells | 1.1 ± 0.8 | 1.5 ± 0.7 | 24.0 ± 10.8 | 0.72 ± 0.25 | 0.03 ± 0.07 | 0.25 ± 0.11 | † | A, V |
| L4 low-threshold spiking | L2/3 cells | Not connected | | | | | | | N/A |
| **L2/3 – L2/3 connections** | | | | | | | | | |
| L2/3 pyramidal | L2/3 pyramidal | 1.0 ± 0.7 | 0.7 ± 0.2 | 15.7 ± 4.5 | 0.61 ± 0.41 | 3.2 ± 7.8 | 0.33 ± 0.18 | 0.10 | F, G |
| | PV+ fast-spiking cell | 0.82±0.49 | 2.32 ± 1.00 | 16.25 ± 5.78 | 0.70 ± 0.14 | 20.0 ± 20.0 | 0.6 ± 0.1* | 0.65 | G, V |
| | PV+ bursting cell | 0.38±0.25 | 2.76 ± 1.05 | 19.2 ± 2.2 | 0.51 ± 0.13 | 13.0 ± 19.8 | 0.5 ± 0.3* | 0.18 | H |
| | Martinotti neuron | 0.25±0.2 | 2.76 ± 1.05 | 19.2 ± 2.2 | 1.91 ± 0.82 | 50 ± 20 | 1.04 ± 0.54 | 0.29 | I, J, P |
| | Neurogliaform cell | 0.39±0.33 | 2.6 ± 0.5 | 17.9 ± 4.0 | 0.83 ± 0.14 | 20 ± 10 | 0.6 ± 0.1* | 0.29 | G, S |
| | CR+ bipolar cell | 1.35±0.62 | 0.64 ± 0.27 | 26.0 ± 8.1 | 0.83 ± 0.14 | 28 ± 22 | 0.39 ± 0.07 | 0.18 | K, N |
| | CR+ multipolar cell | 1.36±0.78 | 1.26 ± 0.53 | 9.5 ± 3.9 | 1.6 ± 0.7 | 50 ± 20 | 0.6 ± 0.1* | 0.20 | M, N |
| | VIP+/CR- cell | 1.35±0.62 | 0.64 ± 0.27 | 26.0 ± 8.1 | 0.83 ± 0.14 | 28 ± 22 | 0.39 ± 0.07 | 0.46 | M, I |
| PV+ fast-spiking cell | L2/3 pyramidal | 0.52±0.45 | 3.5 ± 1.4 | 43.1 ± 10.2 | 0.70 ± 0.15 | 5.1 ± 8.7[5] | 0.46 ± 0.17[5] | 0.60 | G, S |
| | PV+ fast spiking cell | 0.56±0.43 | 1.8 ± 0.6 | 15.8 ± 6.0 | 0.70 ± 0.15 | 5.1 ± 8.7[5] | 0.46 ± 0.17[5] | 0.55 | G, S |
| | others | Not connected | | | | | | | U |
| PV+ bursting | L2/3 pyramidal | 1.21 ± 1.18 | 2.1 ± 1.0 | 22.6 ± 13.7 | 1.27 ± 0.60 | 8.9 ± 11.8[5] | 0.53 ± 0.23[5] | 0.41 | H, O |

| | | | | | | | | | |
|---|---|---|---|---|---|---|---|---|---|
| cell | PV+ fast-spiking cell | 0.77 ± 0.62 | 2.1 ± 1.0 | 22.6 ± 13.7 | 0.86 ± 0.20 | 5.1 ± 8.7[5] | 0.46 ± 0.17[5] | 0.26 | |
| | others | 1.06 ± 0.83 | 2.1 ± 1.0 | 22.6 ± 13.7 | 1.53 ± 0.63 | 8.9 ± 11.8[5] | 0.53 ± 0.23[5] | 0.41 | |
| Martinotti neuron | Martinotti neuron | Not connected | | | | | | | P, T, U |
| | others | 0.29 ± 0.22 | 3.5 ± 1.1 | 13.7 ± 9.9 | 1.80 ± 0.50 | 26.8 ± 26.3[5] | 0.91 ± 0.56[5] | 0.71 | |
| Neurogliaform cell | others | 0.58 ± 0.1 | 53.2 ± 10.8 | 100 ± 19 | 0.51 ± 0.13 | 5.1 ± 8.7[5] | 0.46 ± 0.17[5] | 0.44 | Q, O, R |
| CR+ bipolar cell | L2/3 pyramidal | 0.49 ± 0.49[6] | 5.4 ± 2.2[6] | 56.2 ± 24.1[6] | 1.40 ± 0.50 | 26.8 ± 26.3[5] | 0.91 ± 0.56[5] | 0.11 | L, O, St |
| | PV+ fast-spiking cell | 0.37 ± 0.33[6] | 3.1 ± 2.0[6] | 20.0 ± 12.1[6] | 1.10 ± 0.20 | 26.8 ± 26.3[5] | 0.91 ± 0.56[5] | 0.30 | |
| | CR+ bipolar cell | 0.49±0.56[6] | 4.9 ± 5.4[6] | 33.3 ± 12.0[6] | 1.42 ± 0.20 | 26.8 ± 26.3[5] | 0.91 ± 0.56[5] | 0.32 | |
| | CR+ multipolar cell | 0.49±0.56[6] | 4.9 ± 5.4[6] | 33.3 ± 12.0[6] | 1.33 ± 0.30 | 26.8 ± 26.3[5] | 0.91 ± 0.56[5] | 0.76 | |
| | others | 0.49±0.56[6] | 4.9 ± 5.4[6] | 33.3 ± 12.0[6] | 1.80 ± 0.30 | 26.8 ± 26.3[5] | 0.91 ± 0.56[5] | 0.30 | |
| CR+ multipolar cell | L2/3 pyramidal | 0.49±0.49[6] | 5.4 ± 2.2[6] | 56.2 ± 24.1[6] | 0.7 ± 0.3 | 5.1 ± 8.7[5] | 0.46 ± 0.17[5] | 0.14 | L, O, S |
| | PV+ fast-spiking cell | 0.37±0.33[6] | 3.1 ± 2.0[6] | 20.0 ± 12.1[6] | 1.4 ± 0.4 | 26.8 ± 26.3[5] | 0.91 ± 0.56[5] | 0.18 | |
| | CR+ bipolar cell | 0.49±0.56[6] | 4.9 ± 5.4[6] | 33.3 ± 12.0[6] | 0.98 ± 0.2 | 8.9 ± 11.8[5] | 0.53 ± 0.23[5] | 0.41 | |
| | CR+ multipolar cell | 0.49±0.56[6] | 4.9 ± 5.4[6] | 33.3 ± 12.0[6] | 0.74 ± 0.30 | 5.1 ± 8.7[5] | 0.46 ± 0.17[5] | 0.10 | |
| | others | 0.49±0.56[6] | 4.9 ± 5.4[6] | 33.3 ± 12.0[6] | 1.1 ± 0.4 | 8.9 ± 11.8[5] | 0.53 ± 0.23[5] | 0.50 | |
| VIP+/CR- cell | L2/3 pyramidal | 0.49±0.49[6] | 5.4 ± 2.2[6] | 56.2 ± 24.1[6] | 1.0 ± 0.3 | 8.9 ± 11.8[5] | 0.53 ± 0.23[5] | 0.46[6] | L, O, S |
| | PV+ fast-spiking cell | 0.37±0.33[6] | 3.1 ± 2.0[6] | 20.0 ± 12.1[6] | 1.0 ± 0.3 | 8.9 ± 11.8[5] | 0.53 ± 0.23[5] | 0.38[6] | |
| | others | 0.49±0.56[6] | 4.9 ± 5.4[6] | 33.3 ± 12.0[6] | 1.0 ± 0.3 | 8.9 ± 11.8[5] | 0.53 ± 0.23[5] | 0.38[6] | |

**Supplemental Table 3**

**Features of neural connectivity in the somatosensory (barrel) cortical column.** Pconn, connection probability; PPR, paired pulse ratio. Values are mean ± std.

*Notes:*

† Same parameters are used as in L4-L4 connections, but extended into L2/3. * Values reflect L4-L4 connections. ** References: (A) Beierlein et al 2003; (B) Gil et al 1999; (C) Bruno & Sakmann 2006; (D) Feldmeyer et al 2002; (E) Helmstaedter et al 2008; (F) Sun et al 2006; (F) Feldmeyer et al 2006; (G) Holmgren et al 2003; (H) Blatow et al 2003; (I) Ali 2003; (J) Kapfer et al 2007; (K) Reyes et al 1998; (L) Rozov et al 2001; (M) Porter et al 1998; (N) Caputi et al 2009; (O) Gupta et al 2000; (P) Fino & Yuste 2011; (Q) Wozny & Williams 2011; (R) Tamas et al 2003; (S) Avermann et al 2012; (T) Packer & Yuste 2011; (U) Pfeffer et al 2013; (V) Koelbl et al 2015. [1] Values are taken from A, but scaled according to C.[2] Values are calculated from L4-L4 connections measured in A, based on the difference between L4-L4 synapses and thalamic-L4 synapses reported in B. [3] Values are for thalamus-L4 connection probability. [4] Values are taken from E, as average connection probability between L4 excitatory neurons and L2/3 interneurons.[5] Values are taken from O, based on inhibitory synapse classification. [6] Values are taken from S, as average values for NFS (non-fast-spiking) interneurons.

| | a | b | c | d |
|---|---|---|---|---|
| **L4 neurons** | | | | |
| **Excitatory** | 0.020 ± 0.006 | 0.225 ± 0.014 | -58.0 ± 2.9 | 9.0 ± 1.7 |
| **FS interneuron** | 0.100 ± 0.012 | 0.203 ± 0.015 | -69.7 ± 3.0 | 10.6 ± 1.7 |
| **Non-FS interneuron** | 0.021 ± 0.006 | 0.255 ± 0.028 | -59.9 ± 4.3 | 8.9 ± 1.1 |
| **L2/3 neurons** | | | | |
| **Excitatory** | 0.020 ± 0.006 | 0.225 ± 0.014 | -62.0 ± 4.4 | 11.0 ± 1.7 |
| **PV+ FS** | 0.100 ± 0.011 | 0.213 ± 0.015 | -69.7 ± 3.0 | 12.6 ± 1.8 |
| **PV+ bursting** | 0.021 ± 0.006 | 0.240 ± 0.029 | -54.7 ± 2.7 | 6.9 ± 1.2 |
| **Martinotti** | 0.022 ± 0.006 | 0.225 ± 0.014 | -59.8 ± 4.5 | 8.3 ± 2.1 |
| **Neurogliaform** | 0.021 ± 0.006 | 0.267 ± 0.015 | -85.7 ± 2.7 | 15.9 ± 1.2 |
| **CR+ bipolar** | 0.020 ± 0.006 | 0.260 ± 0.014 | -55.5 ± 3.9 | 8.2 ± 1.5 |
| **CR+ multipolar** | 0.024 ± 0.009 | 0.200 ± 0.012 | -62.1 ± 2.8 | 11.5 ± 1.6 |
| **VIP+/CR- cell** | 0.021 ± 0.006 | 0.230 ± 0.015 | -57.2 ± 4.2 | 8.1 ± 1.6 |

**Supplemental Table 4**

**Table 3.2. Model parameters for different types of neurons in the network.** All values are mean ± std; parameters are normally distributed. FS: Fast-spiking, PV+: Parvalbumin expressing neurons, CR+: Calretinin expressing neurons, VIP+/CR-: Vasopressin expressing Calretinin negative neurons.

| Antibody | Number of image stacks | Average relative difference (%) | Average absolute difference (%) |
|---|---|---|---|
| Anti-NeuN | 13 | 1.24 ± 2.49 | 2.32 ± 1.42 |
| Anti-Parvalbumin (PV) | 7 | -0.55 ± 3.49 | 2.64 ± 2.09 |
| Anti-Calretinin (CR) | 7 | -0.47 ± 3.24 | 2.48 ± 1.91 |
| Anti-Somatostatin (SST) | 6 | 1.19 ± 3.52 | 2.90 ± 2.01 |
| Anti-GAD67 | 7 | 1.68 ± 6.76 | 5.56 ± 3.60 |

**Supplemental Table 5**

**Comparison of automated counting and manual counting results for different antibody staining.** Across all different antibody staining the average relative difference is below 2% when the automated counting results were compared with average results obtained by independent human observers.
